# GEARS: Predicting transcriptional outcomes of novel multi-gene perturbations

**DOI:** 10.1101/2022.07.12.499735

**Authors:** Yusuf Roohani, Kexin Huang, Jure Leskovec

## Abstract

Cellular response to genetic perturbation is central to numerous biomedical applications from identifying genetic interactions involved in cancer to methods for regenerative medicine. However, the combinatorial explosion in the number of possible multi-gene perturbations severely limits experimental interrogation. Here, we present GEARS, a method that can predict transcriptional response to both single and multi-gene perturbations using single-cell RNA-sequencing data from perturbational screens. GEARS is uniquely able to predict outcomes of perturbing combinations consisting of novel genes that were never experimentally perturbed by leveraging geometric deep learning and a knowledge graph of gene-gene relationships. GEARS has higher precision than existing approaches in predicting five distinct genetic interaction subtypes and can identify the strongest interactions more than twice as well as prior approaches. Overall, GEARS can discover novel phenotypic outcomes to multi-gene perturbations and can thus guide the design of perturbational experiments.

## Introduction

The transcriptional response of a cell to genetic perturbation reveals fundamental insights into how the cell functions. It can describe diverse functionality ranging from how gene regulatory machinery helps maintains cellular identity to how modulating gene expression can reverse disease phenotypes [1–3]. This has important implications for biomedical research, especially in the design of more effective and patient-specific therapeutics. For instance, if perturbing the expression of a gene is found to reduce cancer cell proliferation, then a drug targeting that gene would have a significantly higher likelihood of success in clinical trials than one without such target validation [4]. Alternatively, in the multi-gene setting, synergistic gene pairs could be identified that are far more effective in limiting tumor growth when targeted in combination rather than when each gene is targeted individually [5–7]. Knowledge of genetic perturbation outcomes can also dramatically influence the field of stem cell biology and regenerative medicine. Since complex cellular phenotypes are known to be produced by genetic interactions between small sets of genes, these same interactions could also be leveraged to make the precise engineering of cell identity more experimentally tractable [8–12]. While recent improvements in the precision and scale of perturbational screens have enabled scientists to more rapidly sample perturbation outcomes experimentally [8, 13–16], the combinatorial explosion of many possible multi-gene perturbations makes computational approaches indispensable in uncovering outcomes for the vast majority of combinations.

However, existing computational methods for predicting perturbational outcomes present their own limitations. The predominant approach for 1-gene perturbation outcome prediction relies on inferring transcriptional relationships between genes in the form of a network [17–19]. This is limited either by the difficulty in accurately inferring a network from large single-cell gene expression datasets [20] or by the incompleteness of networks derived from existing public databases [21–23]. Moreover, predictive models built using such networks linearly combine the effects of individual perturbations which renders them incapable of predicting non-additive genetic interaction effects [19, 24]. Thus, they cannot predict outcomes for multi-gene perturbations that often exhibit emergent phenotypes such as synergy and epistasis.

More recent work uses deep neural networks trained on data from large perturbational screens to skip the network inference step and directly map genetic relationships into a latent space for multi-gene perturbation outcome prediction [25, 26]. While the use of deep learning enables the prediction of non-additive genetic interactions between combinations of genes, these methods still require that each gene in the combination be experimentally perturbed before the effect of perturbing the combination can be predicted. This is caused by the inability to leverage prior knowledge of genetic relationships, which makes existing models entirely dependent on data from expensive experimental perturbations. For example, the outcome of a 2-gene combinatorial perturbation can only be predicted by such models if both genes have been *seen* experimentally individually perturbed in the training data. By the same reasoning, no 1-gene perturbation outcome can be predicted since that gene would not have been seen experimentally perturbed.

Here, we present GEARS (Graph-Enhanced gene Activation and Repression Simulator), a computational method that integrates deep learning with a knowledge graph of gene-gene relationships to simulate the effects of a genetic perturbation. The incorporation of biological knowledge gives GEARS the unique ability to predict the outcomes of perturbing single genes, or combinations consisting of genes, that were never experimentally perturbed. GEARS uses a new approach of representing each gene and each perturbation with its own multi-dimensional embedding. This allows GEARS to more effectively capture gene-specific heterogeneity and better predict nonlinear interaction effects compared to existing methods. A comprehensive evaluation establishes that GEARS can accurately predict outcomes of 2-gene combinatorial genetic perturbations, significantly outperforming all current approaches. GEARS can predict five different genetic interaction subtypes (synergy, suppression, neomorphism, epistasis and redundancy) and, for four of these, GEARS is twice as accurate as the next best approach in predicting the strongest interactions. GEARS is also able to generalize to new regions of perturbational space by predicting post-perturbation phenotypes that are unlike what was seen during training yet still biologically meaningful. Thus, GEARS can directly impact the design of future perturbational experiments through uncovering a larger region of combinatorial perturbational space than was previously possible using the same experimental data.

## Results

### GEARS combines prior knowledge with deep learning to predict post-perturbation gene expression

GEARS is a deep learning-based model that predicts the gene expression outcome of combinatorially perturbing a set of arbitrarily many genes. The perturbation of each gene in this ‘perturbation set’ is defined as either the activation or repression of the expression of that gene, represented as a signed binary value. GEARS takes as input a vector of expression values that represent an unperturbed cell along with the perturbation set being applied (Figure 1a). The output is the transcriptional state of the cell following the perturbation defined by this set. GEARS is trained using single-cell gene expression data for both unperturbed cells and cells that have undergone known genetic perturbations (Methods).

**Figure 1:**
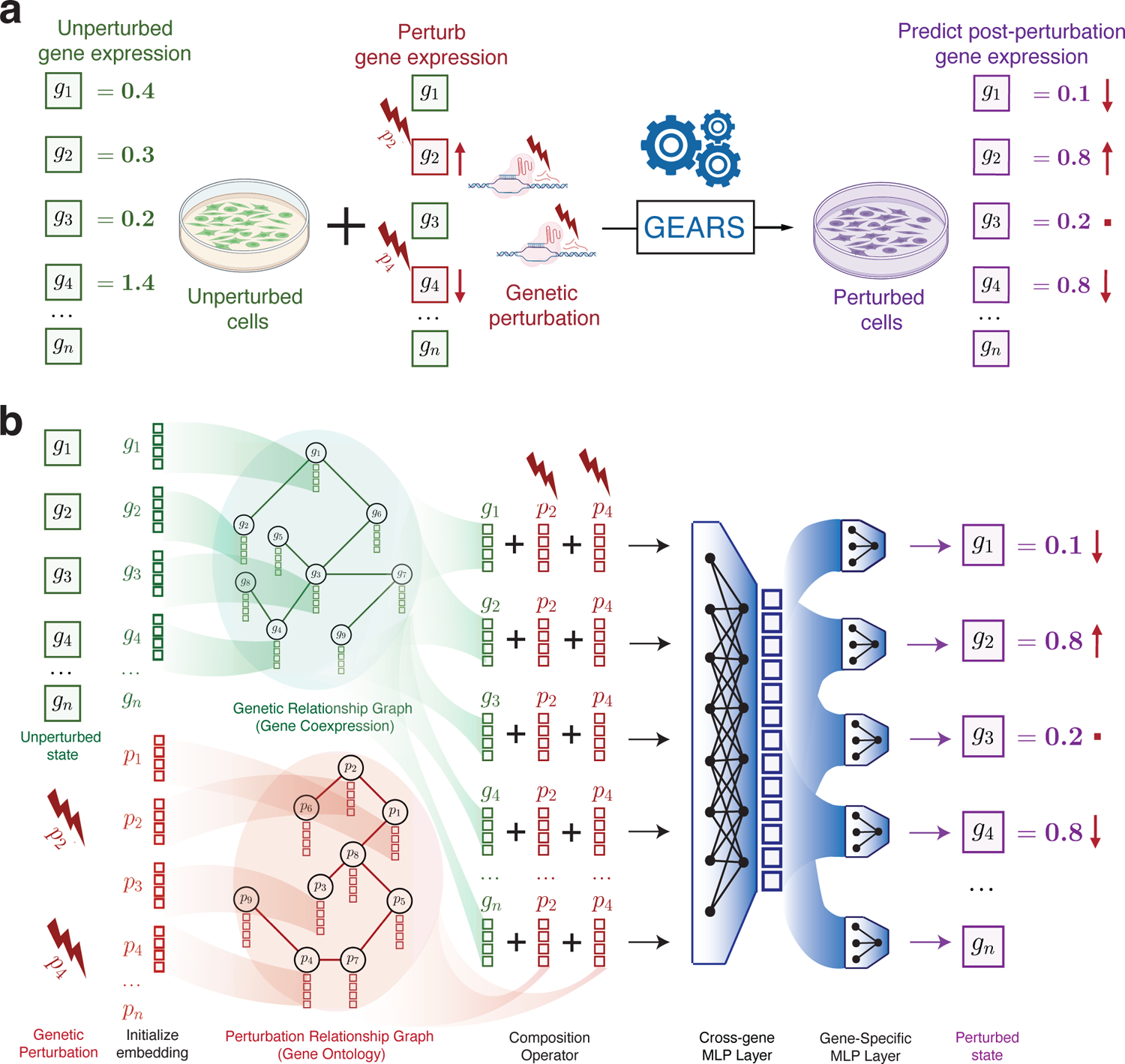
GEARS combines prior knowledge with deep learning to predict post-perturbation gene expression. **(a)** Problem formulation: Given an *n*-dimensional gene expression vector for an unperturbed cell on which a set of genetic perturbations is applied, the goal for GEARS is to predict the gene expression outcome of this perturbation. The perturbation of each gene in the set is defined as either the activation or repression of the expression of that gene. The set can consist of a single gene or multiple genes. **(b)** GEARS model architecture: For each gene in the unperturbed gene expression vector, GEARS initializes a gene embedding vector and a gene perturbation embedding vector. These embedding vectors are assigned as node features in the gene relationship graph and the perturbation relationship graph respectively. A graph neural network is used to combine information between neighbors in each graph. Each resulting gene embedding is summed with the perturbation embedding of each perturbation in the perturbation set. The output is combined across all genes using the cross-gene layer and fed into gene-specific output layers. The final result is post-perturbation gene expression. Crucially, the use of the gene and perturbation relationship graphs allows GEARS to generalize to genes that were never experimentally perturbed during training.

In the case of a multi-gene perturbation, if each gene in the perturbation set has previously been seen experimentally perturbed, then predicting the outcome of that perturbation is trivial unless the interaction effects are not linearly additive. Thus, accurately capturing these non-additive effects is critical for any model that predicts multi-gene perturbational outcomes. GEARS addresses this issue through a new approach of representing each gene and each gene perturbation using its own embedding vector (Figure 1b). By doing so, GEARS more effectively captures the gene-specific heterogeneity of response to perturbation that is responsible for non-linear interaction effects. Each gene’s embedding is sequentially combined with the perturbation embedding of each gene in the perturbation set. This process is independent of the size of the perturbation set making GEARS easily extendable to larger sets. The resulting ‘perturbed’ gene embeddings are combined into a single ‘cross-gene’ embedding vector which captures transcriptome-wide information for each cell. GEARS uses this vector to account for transcriptional effects that were not directly caused by the external perturbation but as a secondary effect of the activation/repression of other genes.

GEARS is uniquely able to extend perturbation outcome prediction to perturbation sets where one or more genes have not been experimentally perturbed. This includes 1-gene perturbations. GEARS does this by not relying entirely on arbitrary encodings to represent each gene perturbation but instead using embeddings that incorporate prior knowledge in the form of gene-gene relationships. The gene co-expression knowledge graph is used as an inductive bias when learning gene embeddings and the same is done using a Gene Ontology-derived knowledge graph for the gene perturbation embeddings (Methods). Here we rely on two biological intuitions: (i) genes that share similar expression patterns should likely respond similarly to external perturbations and (ii) genes that are involved in similar pathways should impact the expression of similar genes upon perturbation (Figure 1b). GEARS is itself independent of the choice and structure of the underlying knowledge graph (Methods). Thus, different knowledge graphs may prove more suitable depending upon the use case and the gene set of interest. GEARS functionalizes this graph-based inductive bias using a graph neural network architecture (Methods) (Figure 1b). Each gene or gene perturbation in the respective input graph is represented as a distinct node and its node-feature vector is set to be its gene embedding or gene perturbation embedding respectively.

### GEARS predicts outcomes for perturbing single genes not seen perturbed during training

In the case of predicting the outcome of 1-gene perturbations, GEARS was evaluated on the perturbation of genes that had been held out at the time of training (Figure 2a). We compare performance with an existing deep learning-based model (CPA) [26] that also learns to represent perturbations in a latent space but models each perturbation using a one-hot encoding and does not use any prior information (Methods). Because no other existing method has this functionality, we also designed two alternative approaches for evaluation of performance. The first (No-Perturb) assumes that the perturbation does not result in any change in gene expression. The second is a linear model which uses the gene co-expression graph to linearly scale and propagate the effect of perturbing a gene (Methods).

**Figure 2:**
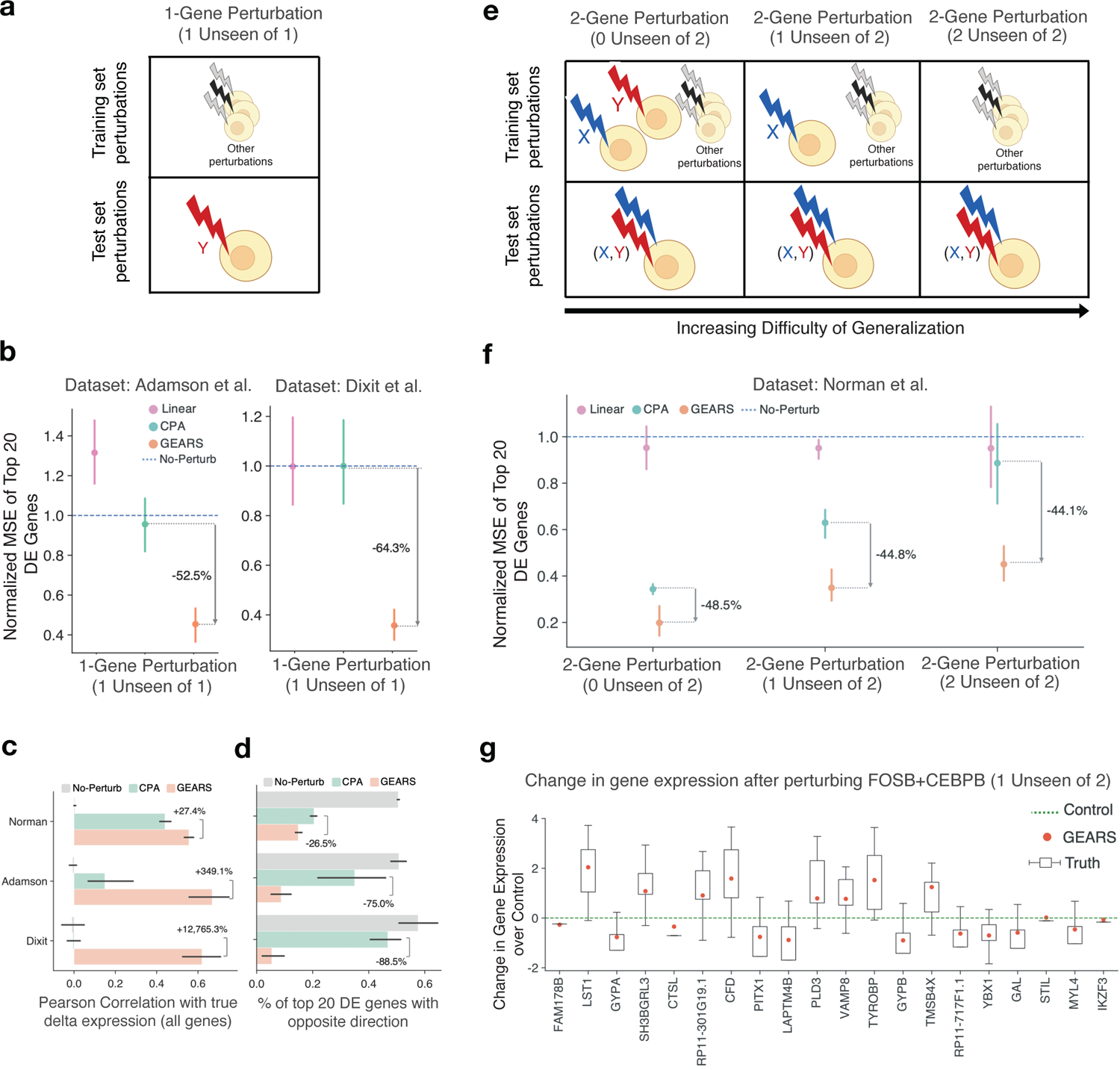
GEARS outperforms alternative approaches in predicting post-perturbation gene expression. **(a)** Train-test data split for 1-gene perturbations. Single gene not seen experimentally perturbed during training is perturbed at the time of testing (1 Unseen of 1) **(b)** GEARS decreases by 50 − 60% the normalized mean squared error (MSE) in predicted post-perturbation gene expression for 1-gene perturbations. For each perturbation, the 20 most differentially expressed genes were considered. MSE is normalized to the no-perturbation case. Perturbation data is from the Adamson et al. dataset [16] and the Dixit et al. dataset [14]. **(c)** GEARS increases the Pearson correlation across all genes by *>* 300% in case of 1-gene perturbations and 26% in case of 2-gene perturbations, as measured between mean predicted post-perturbation differential gene expression over control and mean true post-perturbation differential gene expression over control. **(d)** GEARS increases the percentage of top 20 differentially expressed genes where the predicted post-perturbation expression has opposite direction (activation/inhibition) compared to the ground truth by 75% in case of 1-gene perturbations and by 25% in case of 2-gene perturbation. **(e)** Train-test data split categories for 2-gene perturbations. (i) 2-gene perturbations where both genes in the combination have been seen experimentally perturbed individually at the time of training (0 unseen of 2) and the model then predicts the perturbation result when both genes are perturbed. (ii) 2-gene perturbations where only one (1 unseen of 2) (iii) or none (2 unseen of 2) of the two genes has been seen experimentally perturbed individually at the time of training but at prediction time the model predicts a 2-gene perturbation. **(f)** GEARS increases by 45% the normalized MSE in predicted post-perturbation gene expression for 2-gene perturbations from the Norman et al. dataset [8]. **(g)** GEARS predicts the right trend in gene expression on a gene-by-gene basis. Predicted gene expression across 20 most differentially expressed genes after a combinatorial perturbation (*FOSB+CEBPB*). In this case, only *CEBPB* has been seen experimentally perturbed at the time of training (1 Unseen of 2). The green dotted line corresponds to the mean unperturbed control expression for each gene, the boxes indicate true post-perturbation differential gene expression over control and the red symbol is the mean post-perturbation differential expression predicted by GEARS.

Two different genetic perturbation screens consisting of 87 1-gene perturbations (Adamson et al. [16]) and 24 1-gene perturbations (Dixit et al. [14]) were used for the evaluation. These were run using the Perturb-Seq assay which combines a pooled screen with a single-cell RNA sequencing readout of the entire transcriptome for each cell [14, 16]. Both datasets contained between 50,000 and 90,000 cells with an average of 300-700 cells per perturbation and at least 7,000 unperturbed cells.

GEARS was trained separately on each dataset. A hold-out set of test perturbations was defined for each dataset such that no cell that underwent one of the test perturbations was seen at the time of training. We tested model performance by measuring the mean squared error (MSE) (Figure 2b) and the Pearson correlation (Figure 2c) between the predicted post-perturbation gene expression and the true post-perturbation expression for the held-out set. Since the vast majority of genes do not show significant variation between unperturbed and perturbed states, we restricted our MSE analysis to the harder task of only considering the top 20 most differentially expressed genes. This also makes the evaluation more rigorous since the model cannot trivially predict no perturbation effect for most genes and still achieve a low MSE. GEARS outperforms all baselines significantly on both datasets with an MSE improvement of over 50% (Figure 2b). When looking across all genes using the Pearson correlation, GEARS shows more than three times better performance in the case of both the Adamson and Dixit datasets (Figure 2c). GEARS also shows a clear improvement in capturing the right direction of change in expression following perturbation (Figure 2d) which reflects a more accurate representation of regulatory relationships.

### GEARS predicts multi-gene perturbation outcomes for both previously seen and unseen genes

GEARS predicts outcomes for perturbation sets consisting of multiple genes. However, GEARS was only evaluated on 2-gene perturbations since this was the only combinatorial perturbation data that was publicly available. We used a Perturb-Seq dataset (Norman et al. [8]) containing 131 2-gene perturbations and 105 1-gene perturbations (which included all genes that were perturbed in combination), with 300-700 cells treated with each perturbation. In the case of multigene perturbations, there are multiple categories of generalization which impact the difficulty of the prediction task. Therefore, we defined three such generalization classes when evaluating GEARS on the 2-gene perturbations in the Norman dataset (Figure 2e). The first and simplest case was when the model had seen each of the 2 genes in the combination experimentally perturbed in the training data (2-gene perturbation, 0/2 unseen); the second is when either one of the two individual perturbations had not been seen experimentally perturbed at the time of training (2-gene perturbation, 1/2 unseen) and the third is when both perturbed genes had not been seen experimentally perturbed in the training data (2-gene perturbation, 2/2 unseen) (Supplementary Information Fig 1). GEARS improves performance by approximately 45% across all three levels of generalization (Figure 2f). In fact, even when GEARS was trained with only 1 out of the 2 genes seen experimentally perturbed at the time of training, it was able to perform comparably with the next best performing method that had seen both genes experimentally perturbed.

Model performance was also analyzed on a gene-by-gene basis to make sure that GEARS didn’t overly prioritize some genes over others. In the case of predicting the outcome of perturbing the 2-gene combination *FOSB*+*CEBPB*, GEARS correctly captures both the right trend and the magnitude of perturbation across all 20 differentially expressed genes (Figure 2g) even though one of the genes (*CEBPB*) had not been seen experimentally perturbed during training. GEARS makes accurate predictions in cases of both up and downregulation (e.g. change in the expression of *LST1* and *GYPB*). Similarly, good performance is observed for several other examples across generalization categories (Extended Data Figure2, Supplementary Figure 2).

While the incorporation of knowledge graphs was instrumental in enabling these predictions (Extended Data Figure 4), it also limits GEARS’ ability to predict outcomes for perturbing genes that are both not well connected in this graph and have also not been experimentally perturbed (Methods) (Extended Data Figure 3). GEARS makes use of a Bayesian formulation to overcome this challenge by outputting an uncertainty metric that is inversely correlated with model performance (Supplementary Figure 5). By allowing users to filter out predictions with high uncertainty, this uncertainty metric builds confidence in GEARS’ predictions, especially in the case of perturbation sets containing genes that were not seen experimentally perturbed.

### GEARS can predict new biologically meaningful phenotypes to help uncover the landscape of combinatorial perturbation outcomes

We applied GEARS to the discovery of new phenotypes through predicting the outcomes of all 5,460 pairwise combinatorial perturbations of the 105 genes for which 1-gene post-perturbation expression data was available in the Norman et al. dataset [8] (Figure 3a). GEARS was trained using the post-perturbational gene expression profiles for all 1-gene perturbation outcomes as well as 131 2-gene perturbation outcomes (Figure 3b). The predicted post-perturbation expression captured many distinct phenotypic clusters including all of those previously identified in [8]. Broad trends toward three key lineages of erythroid cells, granulocytes and megakaryocytes were visible (Figure 3c). In addition to these phenotypes, GEARS predicts novel phenotypes that are distinct from those that were observed at the time of training.

**Figure 3:**
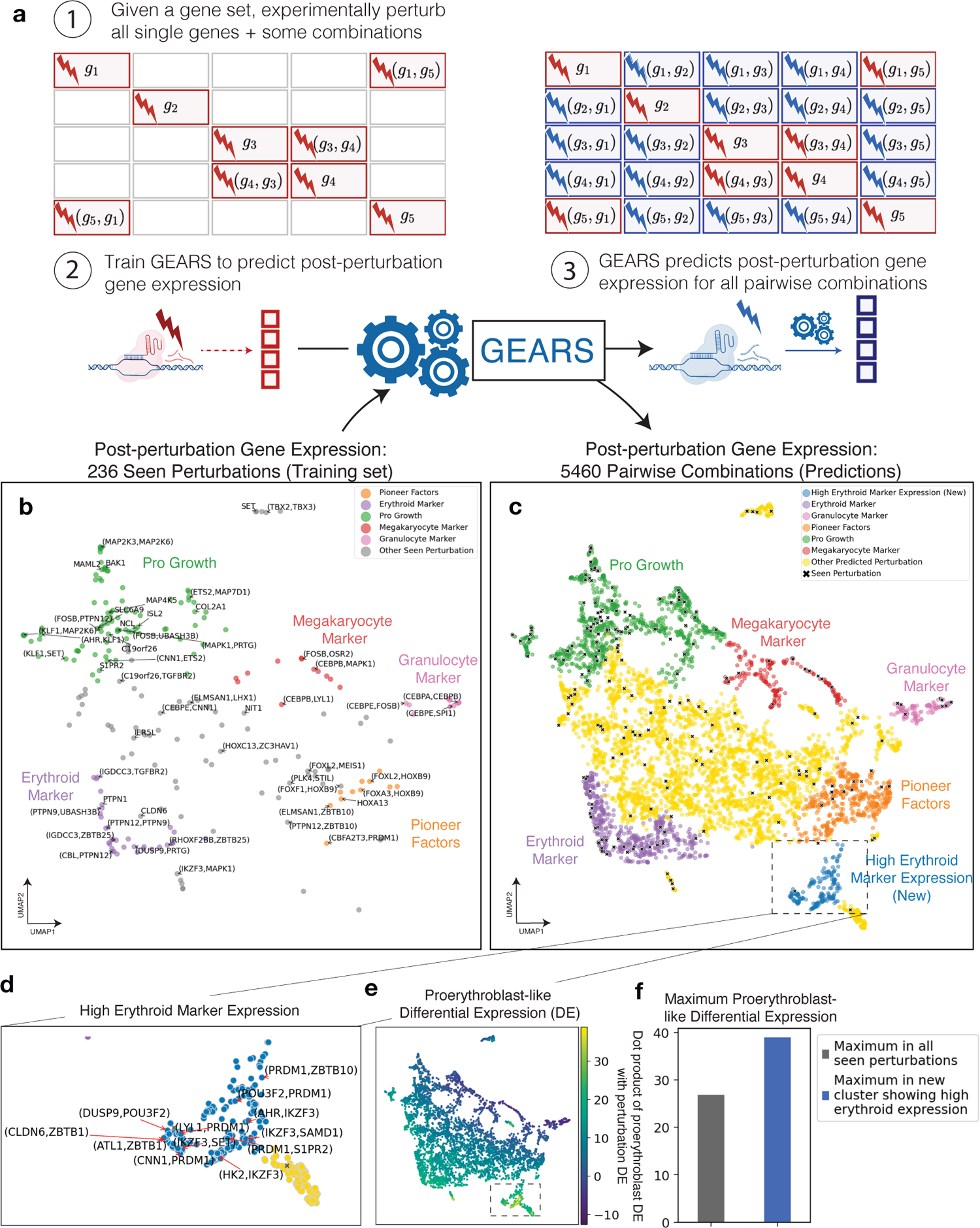
GEARS can predict new biologically meaningful phenotypes to help uncover the landscape of combinatorial perturbation outcomes. **(a)** Workflow for predicting all pairwise genetic interactions for a set of genes (1) Experimentally perturb all single genes in the set and some combinations. (2) GEARS is trained using post-perturbation gene expression for experimentally perturbed genes to predict post-perturbation gene expression for novel perturbations not experimentally perturbed. (3) After training, GEARS predicts post-perturbation gene expression for all pairwise combinations of the gene set. **(b)** Low-dimensional (UMAP) representation of post-perturbation gene expression for experimental perturbations used to train GEARS. Key lineages of erythroid cells, megakaryocytes and granulocytes are visible. The UMAP consists of 105 1-gene perturbations and 131 2-gene perturbations from [8]. A random selection of perturbations is labelled. **(c)** GEARS predicts post-perturbation gene expression for all 5,460 pairwise combinations of the 105 single genes seen experimentally perturbed at the time of training. Low-dimensional (UMAP) representation shows how predicted post-perturbation phenotypes (non-black symbols) are often novel and different from phenotypes seen experimentally (black symbols). Colors indicate Leiden clusters labelled using marker gene expression, following the labeling in [8]. **(d)** GEARS identifies a novel phenotypic cluster of 158 perturbations which displayed significantly higher erythroid marker expression. A random selection of perturbations is labelled. **(e)** Novel cluster identified by GEARS shows differential expression (DE) most similar to proerythroblast-like DE. Color bar measures the dot product between the DE corresponding to the transition from hematopoietic progenitor cells to proerythroblasts (from Tabula Sapiens) and that for the transition from unperturbed controls to each perturbation outcome. **(f)** Maximum proerythroblast-like DE observed for perturbations in the novel cluster is much higher than that observed for any post-perturbation phenotype seen experimentally at the time of training.

For one such cluster showing high erythroid marker expression (containing 158 perturbations including *IKZF3+PRDM1, ATL1+FEV* and *IKZF3+SPI1*), we verified whether the novel phenotype that it represented was biologically meaningful (Figure 3d). Mean differential expression (DE) between unperturbed cells (lymphoblasts) from Norman et al. [8] and each of the genetic perturbation outcomes predicted by GEARS was compared with the DE between hematopoietic progenitor cells and proerythroblast cells (an early stage in the erythroid lineage) in the Tabula Sapiens cell atlas [27]. The goal here was to identify which perturbations produced a change in gene expression that was most similar to that observed in the transition from hematopoietic progenitor cells to an erythroid lineage. The log fold change in expression for differentially expressed genes was used to define a DE *vector* for each of these transitions. Using the dot product between the DE vector for each GEARS-predicted perturbation outcome and the DE vector for proerythroblasts in Tabula Sapiens, we observed that perturbations in the novel cluster showed more similarity to the transition to proerythroblasts than any other perturbation seen at the time of training (Figure 3e, 3f) (Supplementary Information). Thus, this cluster was displaying a phenotype that was not only novel but also biologically meaningful, illustrating how GEARS is able to effectively generalize to new regions of perturbation space. It also highlights how GEARS can be used to discover new experimental routes (perturbations) for engineering cells towards desired phenotypes.

### GEARS predicts non-additive effects of combinatorial perturbation and identifies genetic interaction subtypes

The ability to predict non-additive interaction effects is critical for a multi-gene perturbation model. In the case of a 2-gene perturbation, if the outcomes of perturbing the two genes independently are already known, then a naive model could simply add the perturbational effects to estimate the effect of the combinatorial perturbation (Figure 4a). However, this would not always be accurate since genes are known to interact with one another to produce non-additive genetic interactions (GI) upon perturbation. There are five key GI subtypes: synergy, suppression, neomorphism, redundancy, and epistasis (Methods) (Figure 4b) [8]. For example, two genes that independently cause a minor loss in cell growth could synergistically interact with one another upon combinatorial perturbation to cause cell death. Alternatively, the interaction between two genes could also be epistatic, where one gene dominates the phenotype produced by the combination and masks the effect of the other gene.

**Figure 4:**
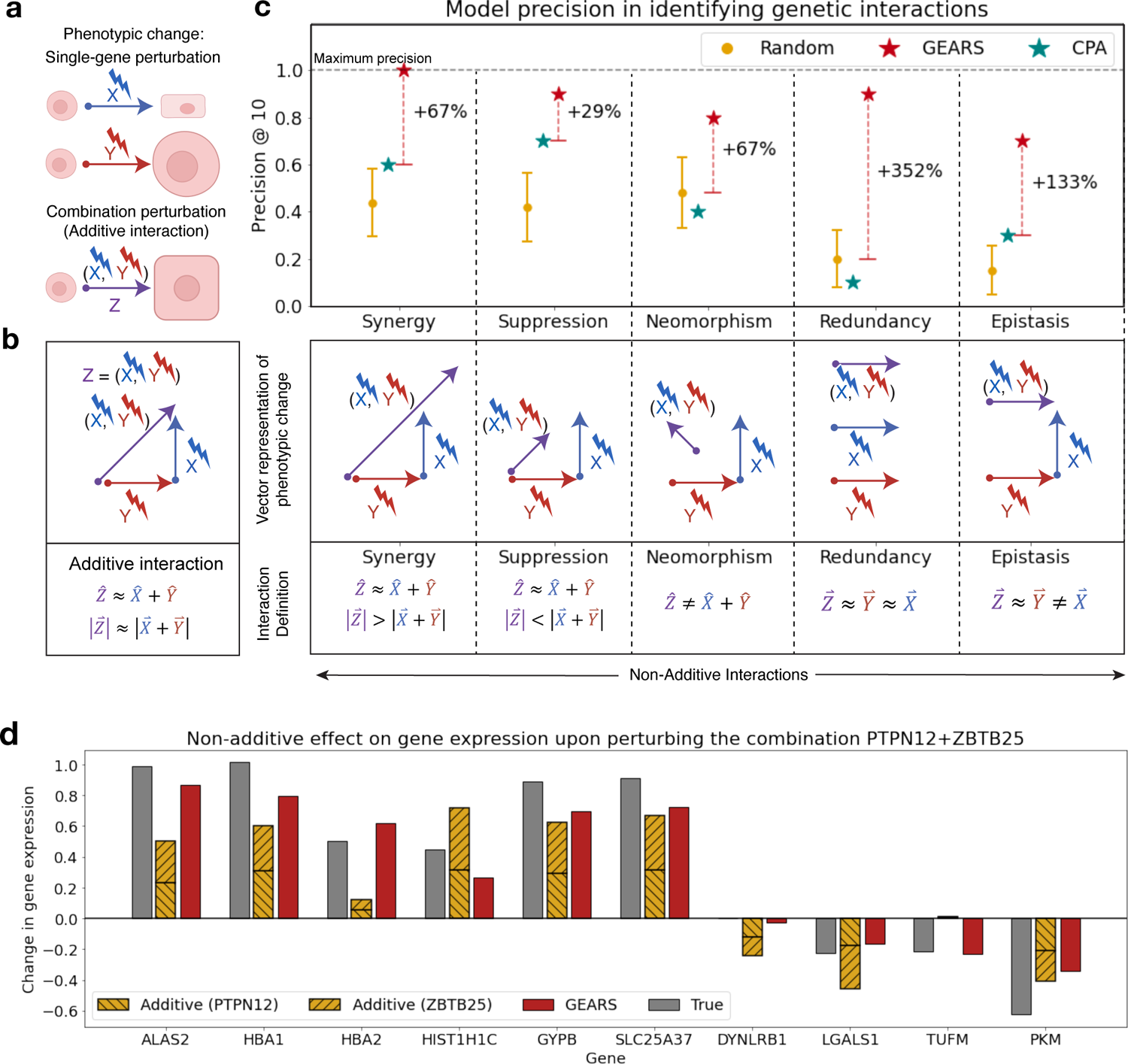
GEARS accurately predicts non-additive combinatorial effects and genetic interaction subtypes. **(a)** Illustration of an additive interaction between two genes upon perturbation. *X* and *Y* represent single-gene perturbations that cause a square-like shape and an increase in the size of the cell respectively. *Z* = (*X, Y*) is a combinatorial perturbation of both genes that results in a larger, square-like cell (i.e. an additive interaction). **(b)** Definition of genetic interactions. Each vector represents the gene expression (phenotypic) change over the unperturbed state caused by a specific perturbation. Under an additive interaction, the true combinatorial phenotype 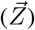 is equivalent to the additive phenotype, i.e. the resultant of the two individual perturbation vectors 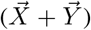. In the case of non-additive interactions, this relationship does not hold true and we see variation in the direction and magnitude of the true combinatorial phenotypic vector as compared to the additive phenotypic vector. Five genetic interaction subtypes are defined. In the case of synergy and suppression, the true combinatorial phenotypic vector is similar in direction to the additive vector but different in magnitude. In the case of neomorphism. the direction is different. Redundancy corresponds to an equivalence between each of the individual perturbations and the combination perturbation. Epistasis occurs when one phenotype masks the effect of the other perturbation in the combination. **(c)** GEARS improves model precision@10 in predicting genetic interactions across all GI subtypes. All 131 2-gene combinations in [8] were ranked using the genetic interaction (GI) score for each GI subtype (Methods). Precision@10 was calculated as the fraction of the top-10 combinations predicted by GEARS for each GI subtype that also showed that GI phenotype based on true post-perturbation expression. The random model corresponds to the result from 1000 random draws. Both GEARS and CPA were trained using a leave-one-out testing approach for each of the 131 combinations. **(d)** GEARS captures different types of non-additive effects at the level of individual genes. Change in gene expression over unperturbed control after perturbing the combination of genes *PTPN12* and *ZBTB25*. The gray bars show the true post-perturbation gene expression change over unperturbed control for a particular gene. The hatched yellow bars show the true post-perturbation gene expression for each of the two single-gene perturbations performed individually. The naive additive model assumes that the effect of the combination is just the sum of the two known single-gene perturbation outcomes. The red bar indicates the prediction made by GEARS. The genes on the y-axis are those with the largest difference between true post-perturbation expression following combinatorial perturbation of *PTPN12* and *ZBTB25* and the additive prediction for that combination.

GIs were defined using metrics (GI scores) that compare observed post-perturbation gene expression with that expected under an additive model. In the case where both genes for each 2-gene combination had been seen experimentally perturbed, GI scores predicted by GEARS showed a very strong correlation to those calculated using true expression (*R*^2^ ≈ 0.5 for all 4 GI scores), much higher than existing methods (e.g. *R*^2^ ≈ 0 in the case of CPA) (Extended Data Figure 5). To simulate a real application of GEARS for recommending experiments, performance metrics were calculated on the top-ranked predictions for each GI subtype. Given the top-10 2-gene combinations predicted to most strongly exhibit a GI subtype phenotype, precision@10 measures what fraction truly exhibits that GI subtype based on experimentally measured post-perturbation gene expression. GEARS increases precision@10 by more than 50% across 4 out of 5 GI subtypes when compared to baseline methods (Figure 4c) with an improvement greater than 100% in the case of redundancy and epistasis. GEARS also shows a doubling in accuracy when directly predicting the set of 10 interactions that are strongest for a GI subtype (Top-10 Accuracy) (Extended Data Figure 6b).

In the novel scenario where one of the two genes in the combination has not been seen perturbed experimentally at the time of training, GEARS also shows significant improvement over baseline approaches. In this case, predictions with high uncertainty were filtered (Methods). When compared to a random baseline, GEARS shows more than a tripling of performance across all interaction types in the case of top-10 accuracy and a doubling of performance in the case of precision@10 for three out of the five GI subtypes (Extended Data Figure 8a, 8b). There was an especially strong performance in the detection of synergy where 72% of all interactions (after filtering for low uncertainty) are correctly detected (Extended Data Figure 8c).

Non-additive interactions can also be evaluated at the level of individual genes. For this, the 20 most non-additively expressed genes were identified for each 2-gene combination. These were the genes where experimentally measured post-perturbation expression deviated most from what was expected under an additive interaction. Based on the MSE for these genes, GEARS is able to capture non-additive effects more than 40% better than existing methods across three out of the five GI subtypes (Extended Data Figure 6a). In the remaining two subtypes, GEARS predictive performance is on par with existing methods. As an example, GEARS was consistently able to predict the correct non-additive effects across almost all of the top 10 non-additively expressed genes following the perturbation of the 2-gene combination *PTPN12*+*ZBTB25* (Figure 4d). These effects were in many different forms; such as synergy in the case of the change in expression of *ALAS2* and *HBA1*, suppression in the case of *HIST1H1C* and neomorphic gene expression in the case of *TUFM*. This was also observed across other examples of combinatorial perturbations belonging to different GI subtypes (Extended Data Figure 9).

### GEARS can effectively search combinatorial perturbation space for novel genetic interactions

GEARS can predict the presence of genetic interactions among all pairwise combinations of a set of genes (Figure 5a). A GI map measuring four different GI scores was generated to simultaneously capture five different GIs: synergy, suppression, neomorphism, redundancy and epistasis. GI scores were calculated using the post-perturbation gene expression predicted for each of the 5.460 pairwise combinatorial perturbations. The GI map reveals a diverse GI landscape where many genes show strong tendencies towards specific GI subtypes (Figure 5b). This effect is most evident in the interactions between functionally related genes which is in line with previous experimental results [13, 14, 28]. For instance, genes involved in early erythroid differentiation pathways (*PTPN12, IKZF3, LHX1*) show a consistent trend of strong synergistic interactions with one another.

**Figure 5:**
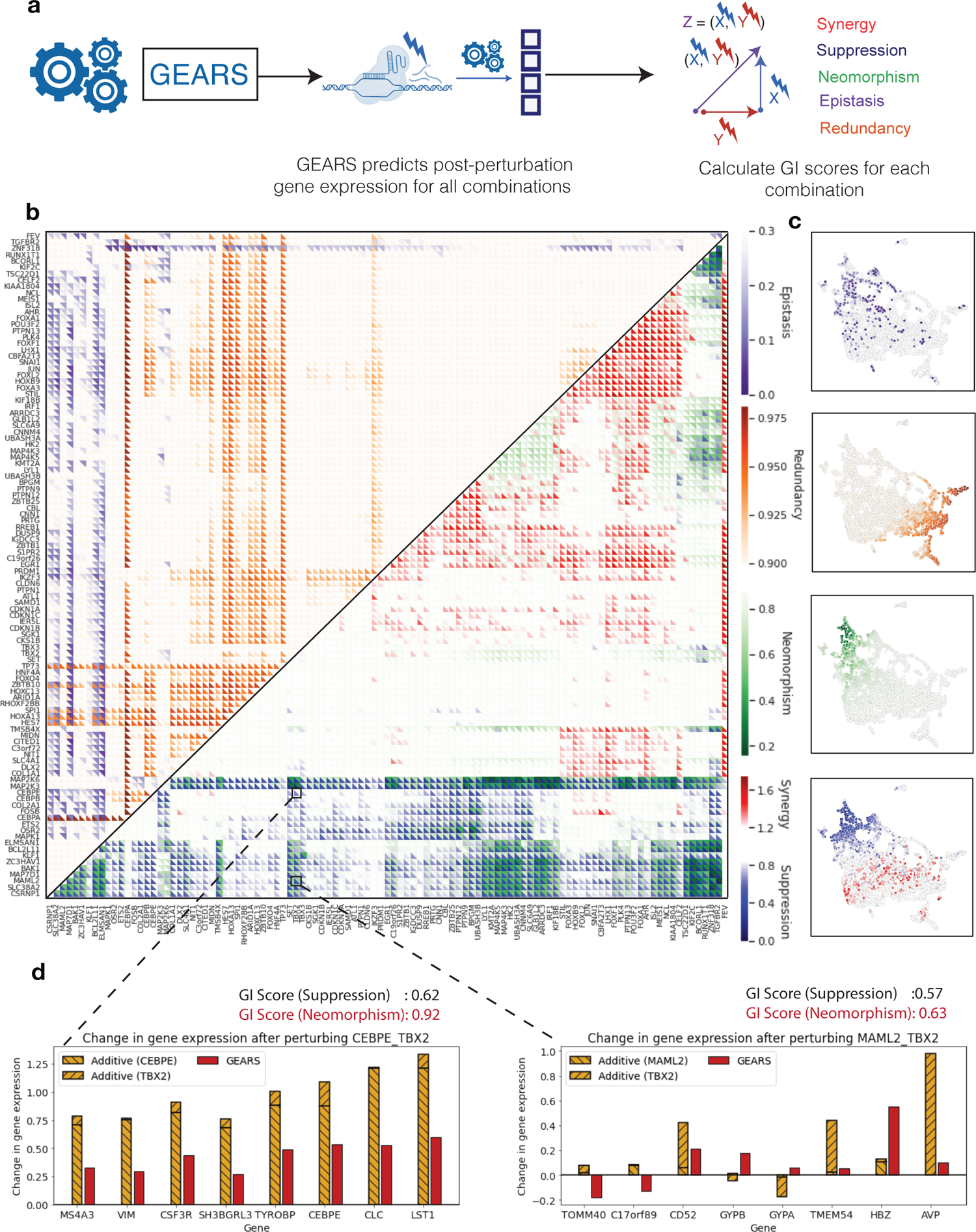
GEARS can search perturbational space for novel genetic interactions of different subtypes. **(a)** Workflow for predicting genetic interaction (GI) scores: First, GEARS predicts the post perturbation gene expression for a given combination. Using this, GI scores are then computed. **(b)** Multi-dimensional GI map generated by GEARS for all pairwise combinations of the 105 single genes perturbed in [8]. For each combination, GEARS predicted GI scores for five different GIs: synergy and suppression (red to blue) measured using the magnitude metric, neomorphism (green) measured using model fit, redundancy (orange) measured using correlation between 1-gene and 2-gene perturbation outcomes and epistasis (purple) measured using the equality of contribution between the two 1-gene perturbations (Methods). The heatmap is clustered by the mangitude metric. **(c)** Genetic interactions are widely distributed across phenotypic space and are often non-overlapping. Low dimensional (UMAP) representation of post-perturbation gene expression for all the perturbations considered in the heatmap (Same UMAP as that in Figure 3(b), 3(c)). UMAPs are colored by the GI scores for each perturbation corresponding to the colorbars used in the heatmap. **(d)** Illustration of how the multi-dimensional GI map can capture significant differences in transcriptional response by accounting for different axes of variability that are not represented in single-dimensional maps. Even though *CEBPE+TBX2* and *MAML2+TBX2* produce the same score for suppression, the GI score for neomorphism indicates that the predicted outcomes are very different. Upon measuring the gene expression changes for the most non-additively expressed genes, *MAML2+TBX2* shows considerable variability in the direction and magnitude of predicted gene expression compared to an additive model. On the other hand, *CEBPE+TBX2* shows a consistent suppressive phenotype.

The uniqueness of this GI map is in how it captures a much broader range of interactions as opposed to conventional GI maps which focus primarily on cell fitness or synergy. For instance, consider the two combinatorial perturbations *CEBPE+TBX2* and *MAML2+TBX2* that would have shown a similar interaction phenotype if only synergy (GI scores: 0.62, 0.57) was being measured. However, GEARS is able to highlight the difference between the two using its measure for neomorphic interactions (Figure 5d), even when they share a common perturbed gene (*TBX2*). The source of this difference is clearly visible when analyzing the most non-additively expressed genes for both perturbations. In the case of *MAML2+TBX2*, GEARS predicts a consistent trend of suppression for these genes without any significant change in direction or scale of expression. However, in the case of *CEBPE+TBX2*, several genes display a change in direction of expression. The non-overlapping nature of different GI subtypes is also clearly visible in the low dimensional UMAP representation of the post-perturbation gene expression for each of the perturbations considered in the GI map (Figure 5c). While neomorphic and redundant interactions tend to cluster in specific regions of this space, epistatic interactions are much more widely distributed. GEARS is further able to identify clusters of activity within each GI subtype. For instance, strongly synergistic combinations tend to produce a similar phenotype for this dataset and distinctly cluster together as opposed to other synergistic combinations.

Finally, the GI map was further expanded to include those combinations where one of the two genes in the combination had not been seen perturbed at the time of training (Extended Data Figure 10). Based on the results of the model evaluation for this harder generalization setting, predictions were only made for synergy and only those under a reasonable threshold of uncertainty were reported. (Extended Data Figure 7). To our knowledge, this is the first example of extending a pairwise GI map beyond those genes that have been seen perturbed individually, opening the door for systematically interrogating much larger regions of perturbational space than was previously possible using the same data.

## Discussion

Predicting transcriptional outcomes of genetic interventions is an important problem in molecular biology with wide-ranging impacts on a number of biomedical research disciplines from regenerative medicine to drug discovery. While recent developments in high throughput perturbational screens have increased both the precision with which genes can be targeted [29, 30] as well as the scale of information generated [15, 31], these experiments remain very costly. Moreover, the combinatorial explosion in multi-gene perturbational space further makes computational methods indispensable for prioritizing which combinations of genes to perturb. However, existing computational approaches face many challenges in fulfilling this potential and are unable to effectively predict multi-gene perturbation outcomes.

We present GEARS, which uses single-cell gene expression data from large perturbational screens to predict outcomes of perturbing novel combinations of genes. GEARS is uniquely able to predict the outcomes of perturbing combinations consisting of genes that have never been perturbed experimentally by leveraging prior knowledge of how genes are interrelated. As CRISPR-based perturbational screens become more ubiquitous for drug discovery, GEARS is uniquely positioned to complement these experiments through inferring an exponentially larger space of multigene perturbation outcomes than existing methods using the same experimental data. Moreover, GEARS can guide the design of new screens by identifying perturbations that would maximize the biological information gained while minimizing experimental cost (Extended Data Figure 3). One constraint in this process is that GEARS must be trained on a particular cell type or a desired experimental condition to make reliable predictions under those same conditions. Building transferability across cell types would help address this issue while also uncovering important insights about how far gene regulatory relationships are shared across cell types.

GEARS is also able to capture gene-specific heterogeneity using a new approach of representing each gene and each gene perturbation with its own multi-dimensional embedding. This allows GEARS to precisely detect the occurrence of non-additive genetic interactions between pairs of genes, especially the strongest interactions for which GEARS is twice as accurate as existing methods. Predicting such emergent behavior is very relevant for discovering tractable routes for engineering cell identity, where cells are guided between transcriptional states that are often significantly different from one another. For instance, GEARS can guide the precise re-engineering of immune cells to prevent exhaustion when targeting cancer [32]. GEARS can also guide the reprogramming of induced pluripotent stem cells to create patient-specifc in-vitro models of disease [33, 34]. Moreover, GEARS is not limited to predicting perturbations that can achieve target states that it has seen at the time of training as it is able to predict novel phenotypes that are biologically meaningful. Overall, this can have significant implications for the field of regenerative medicine. Thus, GEARS can not only impact the discovery of novel small molecules for targeting disease but also push the frontier in the design of the next generation of cell and gene based therapeutics.

## Data availability

All data used for this project has been previously published and the associated citations are referenced in the text.

## Code availability

All code for this project is available at https://github.com/snap-stanford/GEARS

## Methods

### Data preprocessing

The three single-cell RNA-seq datasets used for this study all underwent the same preprocessing. First, each cell was normalized by total counts over all genes and then a log transformation was applied. To reduce the complexity of the prediction problem, we restricted the dataset to only the 5000 most highly varying genes. This is similar to the pre-processing performed by [26] which enabled a more accurate performance comparison (Fig. 2). Since our model requires a gene embedding for every perturbed gene as well, we additionally included any perturbed gene to our dataset that wasn’t already accounted for in the set of most highly varying genes. For the analysis of genetic interactions (Fig. 4), we used the gene set from [8] to ensure consistency in our model’s predictions as compared to the original analysis on the experimental data in [8]. This was generated by identifying all genes in the raw data that had mean UMI value greater than 0.5.

### Overview of GEARS

GEARS considers a perturbation dataset of *N* cells 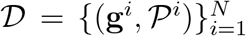, where g^*i*^ ∈ ℝ^*K*^ is the gene expression vector of cell *i* with *K* genes and 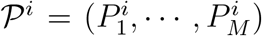 is the set of perturbations of size *M* performed on cell *i*. When *M* = 0, no perturbation is performed and this is the case for an unperturbed cell. Since we are only considering genetic perturbations, each perturbation *P*_*k*_ in the set corresponds to the index of a gene. The goal of GEARS is to learn a function *f* that maps a novel perturbation set 𝒫 to its post-perturbation outcome, which is a gene expression vector g.

Specifically, given a perturbation set 𝒫 = (*P*_1_, ⋯, *P*_*M*_), GEARS first applies an encoder function *f*_pert_ : ℤ → ℝ^*d*^ that maps each genetic perturbation *P* ∈ 𝒫 to a *d*-dimensional gene perturbation embedding. The encoder is a graph neural network (GNN) that operates on the Gene Ontology (GO) graph described later. Another GNN-based encoder function *f*_gene_ : ℤ → ℝ^*d*^ maps each gene into a gene embedding. GEARS then combines the set of perturbation embeddings with each of the gene embeddings using a compositional module to capture genetic interactions. A cross-gene decoder 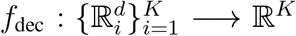 then takes in the set of perturbed gene embeddings and maps them to the post-perturbation gene expression vector. The entire network is trained end-to-end with an auto-focus direction-aware loss.

### Gene co-expression GNN encoder

GEARS first obtains a faithful representation for each gene that captures co-expression patterns in the cell, these are assumed to act together across perturbations. We observe that the relative heterogeneity of perturbational response is high for each gene, suggesting that the model should assign capacity to capture this heterogeneity. Thus, instead of representing each gene as a scalar, GEARS represents each gene *u* ∈ ℤ as a learnable embedding **x**^gene^ ∈ ℝ^*d*^.

To enable the gene embeddings to reflect these co-expression relationships, we apply a GNN on a constructed gene co-expression graph 𝒢_gene_. Particularly, nodes in 𝒢_gene_ are genes and edges link co-expressed genes. GEARS calculates Pearson correlation *ρ*_*u,v*_ among genes *u, v* in the training dataset. For each gene *u*, we connect it to the top *H*_gene_ genes that have highest *ρ*_*u,v*_ and are above some threshold *δ*. Next, we apply a GNN parameterized by *θ*_*g*_ that augments every gene *u*’s embedding 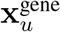, by integrating information from the embeddings of its co-expressed genes (i.e. neighboring genes in 𝒢_gene_): 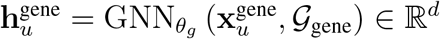.

### Injecting perturbation structure with gene ontology GNN

GEARS predicts the outcome of perturbing genes never seen perturbed before through leveraging a key intuition that perturbation responses are extremely similar for genes that are involved in the same pathways. This observation suggests that we could build a representation of novel gene perturbations by learning from a composition of previously seen gene perturbations that share the same pathways as recorded in the Gene Ontology graph [35] 𝒢_GO_. GEARS leverages this prior knowledge and injects it into the model through a GNN encoder.

More specifically, we first construct a gene perturbation similarity graph 𝒢_pert_ based on the Gene Ontology graph 𝒢_GO_. 𝒢_GO_ is a bipartite graph where an edge links a gene to a pathway GO term. We denote 𝒩_*u*_ as the set of pathways for a gene *u*. We compute Jaccard index between a pair of gene *u, v* as 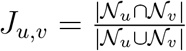. It measures the fraction of shared pathways between the two genes. For each gene u, we then select the top *H*_pert_ gene *v* with the highest *J*_*u,v*_ to construct 𝒢_pert_. Next, we initialize all possible gene perturbations (*P*_1_, ⋯, *P*_*K*_) with learnable embeddings 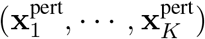. We then feed them into a GNN parameterized by *θ*_*p*_ to augment every perturbation *v*’s embedding 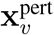 by integrating information from perturbation embeddings that share similar pathways (i.e. neighboring perturbations in 𝒢_pert_): 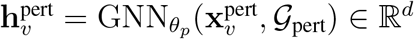.

### Modeling combinatorial perturbation across genes

During training, GEARS maps each gene to a perturbation embedding using the gene ontology GNN above. Given a perturbation set 𝒫 = (*P*_1_, ⋯, *P*_*M*_), GEARS looks up the perturbation embedding of each element of that set 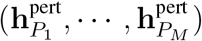. To model the genetic interactions among multiple perturbations, we use the ‘sum’ compositional operator followed by a multi-layer perceptron 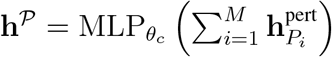. The ‘sum’ operator allows extendability to perturbations of any size. Thus, each perturbation embedding from 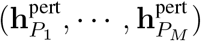 is applied to every gene embedding to obtain a post-perturbation gene embedding. For gene *u*, we have 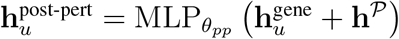.

### Cross-gene gene-specific decoder

Following the application of the perturbations in the embedding space, GEARS maps the post-perturbation gene embedding to its corresponding post-perturbation gene expression vector. Since each gene has its own perturbation pattern, for every gene *u*, we apply a gene-specific linear layer parameterized by **w**_*u*_ ∈ ℝ^*d*^, *b*_*u*_ ∈ ℝ to map it to a scalar of perturbation gene expression effect 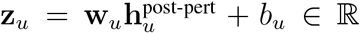. We then concatenate the individual effect to a single perturbation effect vector **z** ∈ ℝ^*K*^ for the cell. Since the perturbational effect on a gene can incur secondary effects on other genes, we wanted to use the transcriptome-wide ‘cross-gene’ information for the cell when predicting final gene expression for each gene. Thus, we added an additional MLP that generates a cross-gene embedding for the cell 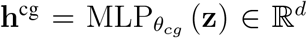. Conditioned on this cross-gene state, for every gene *u*, a gene-specific decoder parameterized by 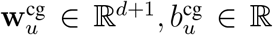 augments **z**_*u*_ to 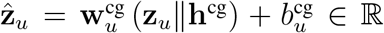. Finally, the predicted perturbation effect vector 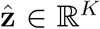 is added to the gene expression of a randomly sampled unperturbed control cell to arrive at the predicted post-perturbation gene expression vector for that cell 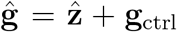. Thus, GEARS learns to predict the change in gene expression over control following perturbation instead of the absolute post-perturbation expression. This allows it to avoid allocating model capacity on learning basal gene expression and instead focus on learning perturbation effects.

### Autofocus direction-aware loss

GEARS optimizes model parameters to fit the predicted ĝ post-perturbation gene expression to true post-perturbation gene expression g using stochastic gradient descent. We observed that majority of genes incur minimal perturbational effects. Since we are most interested in the differentially expressed genes, we designed an autofocus loss that automatically give a higher weight to differentially expressed genes by elevating the exponent of the error. Particularly, given a minibatch of *T* perturbations, where each perturbation *k* has *T*_*k*_ cells, and each cell has *K* genes with predicted post-perturbation gene expression ĝ and true expression g, the loss is defined as:

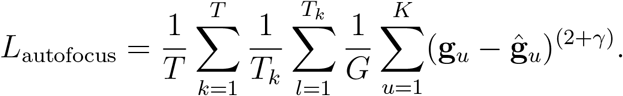

In addition to the absolute value of perturbation effect, the direction of change in expression compared to control is also important since it captures whether the perturbation activates or inhibits a gene. A standard loss is insensitive to this directionality. To address this, GEARS incorporates an additional direction-aware loss:

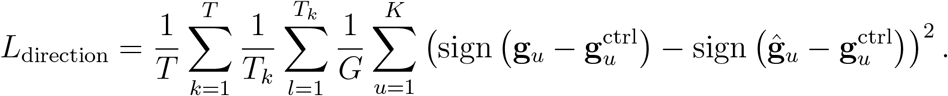

The prediction loss function is *L* = *L*_autofocus_ + *λL*_direction_, where *λ* adjusts the weight for the directionality loss.

### Uncertainty

GEARS generates an uncertainty score to measure the confidence of model prediction on a novel perturbation. GEARS fixes a Gaussian likelihood 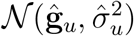 to model the post-perturbation gene expression value for gene *u* under perturbation 𝒫, where ĝ_*u*_ is the predicted post-perturbation scalar, and 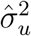 is the variance [36]. We add an additional gene-specific layer to predict the log-variance term 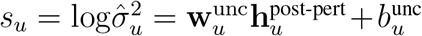 for each gene u and learn it through a modified bayesian neural network loss [36]:

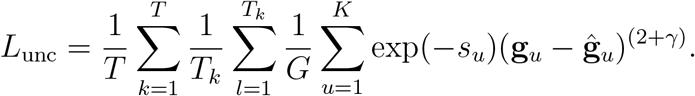

Mechanistically speaking, the loss encourages log-variance to be large when the error is large. Thus, the log-variance is learned to be a proxy of model uncertainty. If the uncertainty score is desired at the time of inference, GEARS simply needs to update the prediction loss function *L* by adding the uncertainty loss *L*_unc_.

### Hyperparameters

We use HyperBand [37] on the validation set of a fixed split of the Norman dataset to find the best hyperparameters. The same set of hyperparameters are then used across all datasets and multiple splits. The set of ranges for the hyperparameters include: GNN architecture – {graph convolutional network (GCN) [38], graph attention network (GAT) [39], simplifying graph convolutional network (SGC) [40]}; GNN layer size – {1, 2, 3}; hidden size *d* – {32, 64, 128}; autofocus loss coefficient γ – {2, 4}; direction loss regularization term *λ* – {1, 0.1, 0.01}; the number of top similar genes in the co-expression network *H*_pert_ – {3, 5, 10, 20}; the number of top similar genes in the perturbation network *H*_gene_ – {3, 5, 10, 20}; correlation threshold for co-expression network *δ* - {0.4, 0.8}; learning rate – {1e-2, 1e-3, 1e-4}; batch size – {32, 64, 128}. Since we have a large set of hyperparameters, for a more efficient selection, we apply HyperBand on different groups of hyperparameters where each group has a small set of hyperparameters while fixing the rest. The final set of hyperparameters are the following: GNN architecture - SGC; GNN layer - 1; hidden size - 64; γ, - 2; *λ* − 0.1; *H*_pert_ - 20; *H*_gene_ - 5; *δ* = 0.4; learning rate - 1e-3; batch size - 32.

### Using a graph to represent prior knowledge

GEARS does not require a specific representation of prior knowledge about gene-gene relationships. For capturing similarities between the gene embeddings we chose to use the gene coexpression graph. For the gene perturbation embeddings, we used the Gene Ontology graph which was generated by adding weighted edges between genes that shared a significant number of GO terms. The generation procedures for both graphs were described previously. We also experimented with a few different networks to use in place of the Gene Ontology network including a protein-protein interaction network [21], a gene coessentiality network [41] or the gene co-expression network described above. We decided to proceed with the Gene Ontology network because it had the best coverage over the gene set that we were interested in, produced very good predictive performance and was the most general-purpose for application to future tasks.

### Model evaluation for predicting overall gene expression

For predicting overall gene expression, we used the mean square error between the model predictions and the true post-perturbation gene expression for perturbations held out in the test set. Generally, it is quite expensive (if not impossible) to perturb all genes or all combinations of genes when running a perturbational screen. This makes it very useful to be able to computationally predict perturbational response to perturbations that were not seen at the time of training.

To simulate this real world scenario, we constructed a data split to account for all possibilities for single-gene and 2-gene perturbations. In the case of 2-gene combinations, there are three possible types of perturbations from a data split perspective: (1) combinatorial perturbations where both single-gene perturbations in the combination have been seen perturbed individually at the time of training (2-gene perturbation, 0/2 unseen); (2) those where only one of the two single-gene perturbations have been seen perturbed individually at the time of training (1/2 unseen) or perturbations where neither of the two single-gene perturbations have been seen perturbed individually at the time of training (2/2) unseen. In the case of single-gene perturbations, there is only one category which is simply those perturbations that were not seen at the time of training (1-gene perturbation 1/1 unseen).

To generate a data split for the Norman et al. dataset which contained both single and 2-gene perturbations [8], we first randomly sample *K*_*G*_% from the gene list and consider them as the gene set that is seen at the time of training. Thus, all single-gene perturbations with genes belonging to this set are used for training. The rest of the genes (1-*K*_*G*_)% are used as the unseen gene set and the corresponding single-gene perturbations are used for testing. Next, within the 2-gene combination perturbations, in the case when both individual perturbations are in the seen set (0 unseen of 2), we randomly sample *K*_*C*_% of them as training perturbations and the rest (1-*K*_*C*_)% are held out in the test set. For the other 2 categories: 1/2 unseen and 2/2 unseen, we simply hold out all 2-gene combinations where at least one of the individual genes being perturbed in that combination is in the unseen set. See Extended Figure 1 for an illustration. In our study, we set *K*_*G*_ = 75, *K*_*C*_ = 75 to obtain the train+validation and test set and then in the train+valid set, we run *K*_*G*_ = 90, *K*_*C*_ = 90 to obtain the train and validation set. In the case of datasets containting only single-gene perturbations (Adamson et al. [16], Dixit et al. [14]), we only test performance on single-gene perturbations which were not seen perturbed at the time of training (single unseen).

### Baseline models

The following baseline models were used for comparing model performance:

1. **No perturbation model**: This model simply predicts that there was no effect of performing a perturbation and that the unperturbed cell state is the same as the post-perturbed one.
2. **Linear model**: This model uses the gene coexpression graph to learn weights between all the genes. When a perturbation is applied to a specific gene, it propagates the effect of that perturbation to its neighbors through its edges which linearly scale the magnitude of that perturbation. Those neighbors in turn will further propagate the effect of that perturbation to their own neighbors. We allowed perturbations to propagate this way upto 3 hops away from the site of the original perturbation. Let *E* represent the adjacency matrix of the weighted gene coexpression graph and *θ* represent a genetic perturbation vector. *θ* is *n*-dimensional vector where n is the number of genes. It has a value of zeros at every position except at the indices of genes where perturbations are being applied, where it is either +1 or − 1. Let *d* = 3 be the number of hops. Then, the change in gene expression **x**_*θ*_ caused by a perturbation *θ* under the linear model would be:

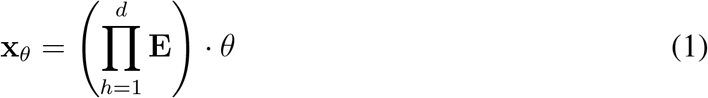
3. **Compositional Perturbation Autoencoder (CPA)** [26]: This model uses an adversarial autoencoder with no other prior information to predict the effect of applying a specific perturbation to a given unperturbed cell.

### Measuring genetic interaction scores

For identifying and categorizing genetic interactions we followed the definitions and metrics defined in Norman et al. [8]. They defined the following types of GIs: additive, epistatic, neomorphic, potentiation, redundant, suppressive, synergy (similar/dissimilar). The authors make a distinction between synergistic combinations based on the similarity of the combining single-gene perturbations. We did not include this division because our focus was on evaluating predictions for combinatorial perturbations. We also did not include ‘potentiation’ as a separate category and instead grouped it under synergy. This is because it was defined as the combined interaction of high synergy and epistasis and we evaluated those GIs individually. ‘Additive’ interactions (or the no-GI class), which are defined as the complement of seeing either synergy or suppression are only included in Extended Data Figures 4, 5.

Norman et al. [8] defined metrics (GI scores) for identifying GIs using a linear model of the combinatorial perturbation effect. Let **g**^*i*^ ∈ ℝ^*K*^ be the post-perturbation gene expression vector of a cell *i* with *K* genes. Let *C*_*k*_ be the set of cells under perturbation *k*, where |*C*_*k*_| = *T*_*k*_. The first step is to compute the average post-perturbation gene expression 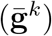 for each of the two combining genes *a, b* perturbed singly as well as in combination (*a* + *b*):

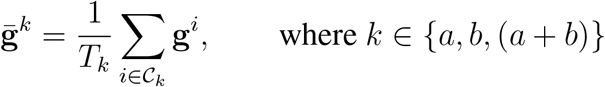

Then the change over mean expression in unperturbed control cells 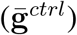 is computed as:

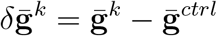

And it is used to fit the following linear model:

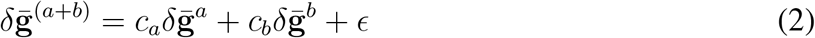

Here *ϵ* captures the error in the model fit. Following the procedure in Norman et al [8], the model was fit using robust regression with a Theil-Sen estimator (fit on 10,000 random subsamples of 1,000 genes at a time) Using the values of the coefficients, the following metrics (or GI scores) were defined shown below. To simplify the notation we write 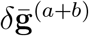 as **ab**, 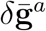 as **a** and 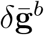 as **b**.

|  | Metric (GI score) | Definition | Relevant GI |
| --- | --- | --- | --- |
| 1 | Magnitude | $\sqrt{c_a^2 + c_b^2}$ | Synergy, Suppression, Additivity |
| 2 | Similarity of (single/double) transcriptional profiles | $corr([\mathbf{a}, \mathbf{b}], \mathbf{ab})$ | Redundancy |
| 3 | Model fit | $corr(c_a \mathbf{a} + c_b \mathbf{b}, \mathbf{ab})$ | Neomorphism |
| 4 | Equality of contribution | $\frac{\min(dcor(\mathbf{a}, \mathbf{ab}), dcor(\mathbf{b}, \mathbf{ab}))}{\max(dcor(\mathbf{a}, \mathbf{ab}), dcor(\mathbf{b}, \mathbf{ab}))}$ | Epistasis |

Here, *corr* refers to a distance correlation and the square brackets represent the concatenation operation. When predicting a GI score, first the mean post perturbation expression vectors are predicted for both the combination perturbation and the single-gene perturbations 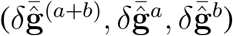. These are then used to estimate the relevant parameters such as in (2). When calculating the true value for the GI score, the same procedure is performed with true post perturbation gene expression vectors 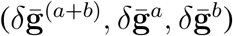.

### Identifying genetic interaction subtypes

For each defined GI subtype q, the authors in [8] defined a set of 2-gene combinatorial perturbations **S**_*q*_ as expressing that type of interaction. However, they did not explicitly state the GI score thresholds used to define these sets. To estimate these thresholds, we first computed the relevant GI score for every element belonging to a given GI subtype set **S**_*q*_ using true post-perturbation gene expression. We then estimated the minimum score in case of a lower bounded condition and the maximum score in case of an upper bounded condition and used this as the score threshold *τ*_*q*_ for each GI subtype *q*. These thresholds are also visualized as colored horizontal lines in Extended Data Fig 5. The result was the following conditions for labeling an interaction as belonging to a specific GI subtype. Overall, no GI subtype set accounted for more than 50% of all 131 combinations being tested.

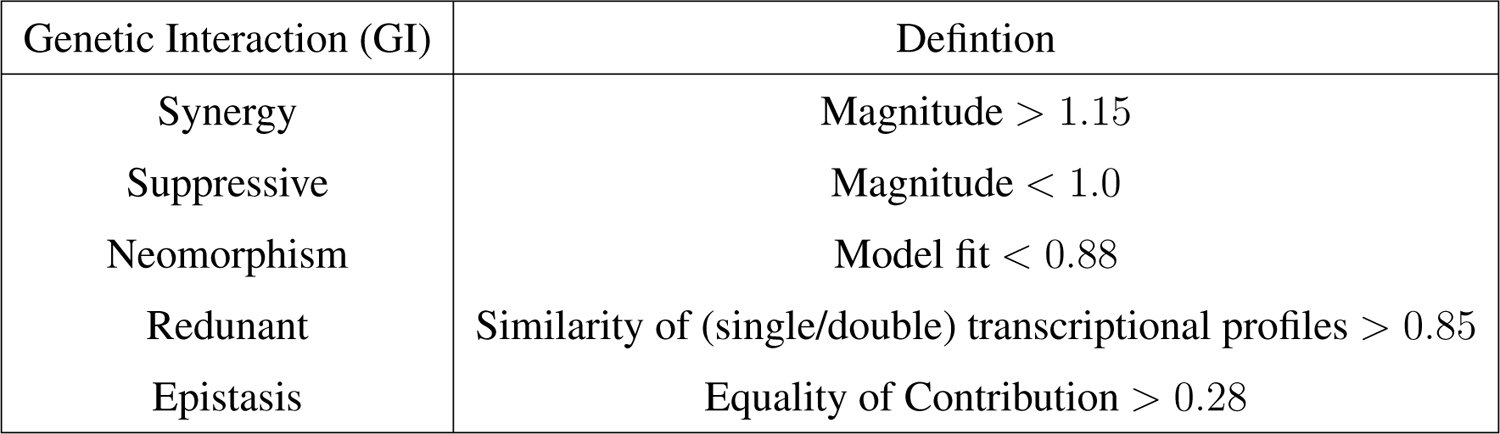

| Genetic Interaction (GI) | Defintion |
| --- | --- |
| Synergy | Magnitude $> 1.15$ |
| Suppressive | Magnitude $< 1.0$ |
| Neomorphism | Model fit $< 0.88$ |
| Redunant | Similarity of (single/double) transcriptional profiles $> 0.85$ |
| Epistasis | Equality of Contribution $> 0.28$ |

### Model evaluation for predicting genetic interaction

We evaluated GEARS’s ability to correctly predict different GI subtypes. A leave-one-out testing procedure was followed for this analysis. For every combinatorial perturbation experimentally tested in Norman et al. [8], we trained GEARS from scratch while only holding out that specific interaction in the test set. Thus, we trained 131 different models. We performed the same procedure with the deep learning-based baseline model CPA [26].

Once each model was trained, we computed all the GI scores (Table 1) for the perturbation that was held out in the test set. Using thresholds from Table 2, we identified whether a specific GI was predicted to exhibit a specific GI subtype. The same procedure was also performed using true post perturbation gene expression. The performance of GEARS in predicting each GI subtype was evaluated using the following metrics

- **Precision**: The fraction of combinatorial perturbations predicted to show a specific GI subtype that were also identified to do so based on true post-perturbation expression (Extended Data Figure 5,6). Let **Ŝ**_*q*_ be the set of perturbations predicted to show a specific GI subtype and **S**_*q*_ be the perturbations that truly show that GI subtype.

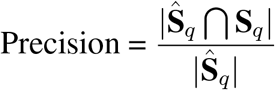
- **Recall**: The fraction of combinatorial perturbations that were identified as showing a specific GI subtype based on true post perturbation gene expression that were also predicted to do so by the model being evaluated (Extended Data Figure 5,6).

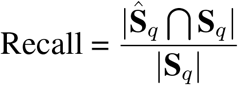
- **Precision@10**: Of the 10 combinatorial perturbations predicted to have the highest GI score for a given GI subtype, precision@10 refers to the fraction that were truly identified as belonging to that GI subtype using true post-perturbation expression. For example, the ten combinatorial perturbations with the highest score for magnitude were used to evaluate precision@10 for synergistic interactions while those with the lowest were used to do the same for suppressive interactions. Let 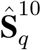 be the set of 10 combinatorial perturbations predicted by a model to have the highest GI score for a given GI subtype. Here |**S**_*q*_| ≥ 10.

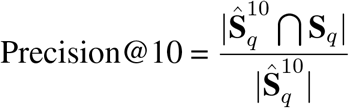

In practice, it is more common for scientists to choose a handful of promising combinations to test experimentally as opposed to exhaustively testing all likely combinations. Thus, by focussing on the model’s ability to correctly rank the most likely genetic interactions, precision@10 captures the success probability of follow-on experiments that aim to validate model predictions. We compared our performance to a random baseline by drawing 1000 random sets of 10 combinations from this set and plotting their mean and standard deviation as a null model. This set of 131 combination perturbations was slightly biased towards the presence of an interaction, thus the random baseline helps to put our predictive performance in context. We did not use the naïve baseline that assumed that the combination perturbation effect would be a simple sum of the single-gene perturbation effects, because this would trivially result in the same GI score for all combinations.
- **Top-10 Accuracy**: Of the 10 combinatorial perturbations predicted to have the highest GI score for a given GI subtype, top 10 Accuracy refers to the fraction that were also identified as being part of the 10 combinatorial perturbations identified to have the highest GI score using true post-perturbation expression. Thus, top-10 accuracy is more robust to biases in the dataset towards oversampling genetic interactions but it is also a more conservative metric for measuring performance. Let 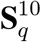 be the set of 10 combinatorial perturbations identified to have the highest GI score for a given GI subtype as measured using true post-perturbation gene expression.

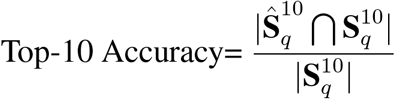

### Model evaluation for predicting non-additive effects

The GI scores defined above [8] consider the expression values of all genes when calculating the score. Often, it is only the expression of a few genes that manifests an interaction phenotype or a non-additive effect. To focus our analysis on these interacting genes, we measured how many genes post-perturbation were expressed in a manner that was very different from a simple additive effect. We first defined a naïve additive model that simply added together the effects of the individual single gene perturbations. As defined previously, if 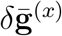 represents the mean change in expression over unperturbed control when perturbing gene *x*, then the naive additive model predicts that the effect of perturbing the combination of genes (a + b) would result in the following effect:

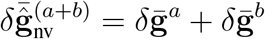

We used this naive sum to sort genes by how far their true post-perturbation expression under a combination perturbation deviated from this naïve prediction (Fig. 3a).

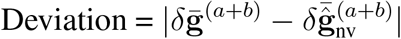

We then measured the mean squared error in predicting the top 20 with the highest deviation across all combination perturbations. The final results were categorized by GI type (Figure 3b).

### Selecting predictions with low uncertainty

GEARS is able to predict an uncertainty value 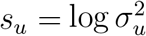 for each gene *u*. To generate a transcriptome-level uncertainty value, we simply took the mean across all model-predicted uncertainty values for all genes. So, for some cell *i*, we estimated its uncertainty value as the following:

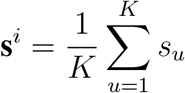

To allow comparison of this uncertainty value across different models, we performed z-score normalization using the mean and standard deviations of the predicted uncertainty values for all the data used to train that model. If 𝒞_*tr*_ are the cells in the training data, we first calculate the mean *µ*_*tr*_ and standard deviation *σ*_*tr*_ of the set of uncertainty values {**s**^*i*^ : ∀*i* ∈ 𝒞_*tr*_}.

We can then z-score normalize the uncertainty values for any cell *j* across different trained models as follows:

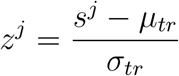

## Extended Data

**Extended Data Fig. 1:**
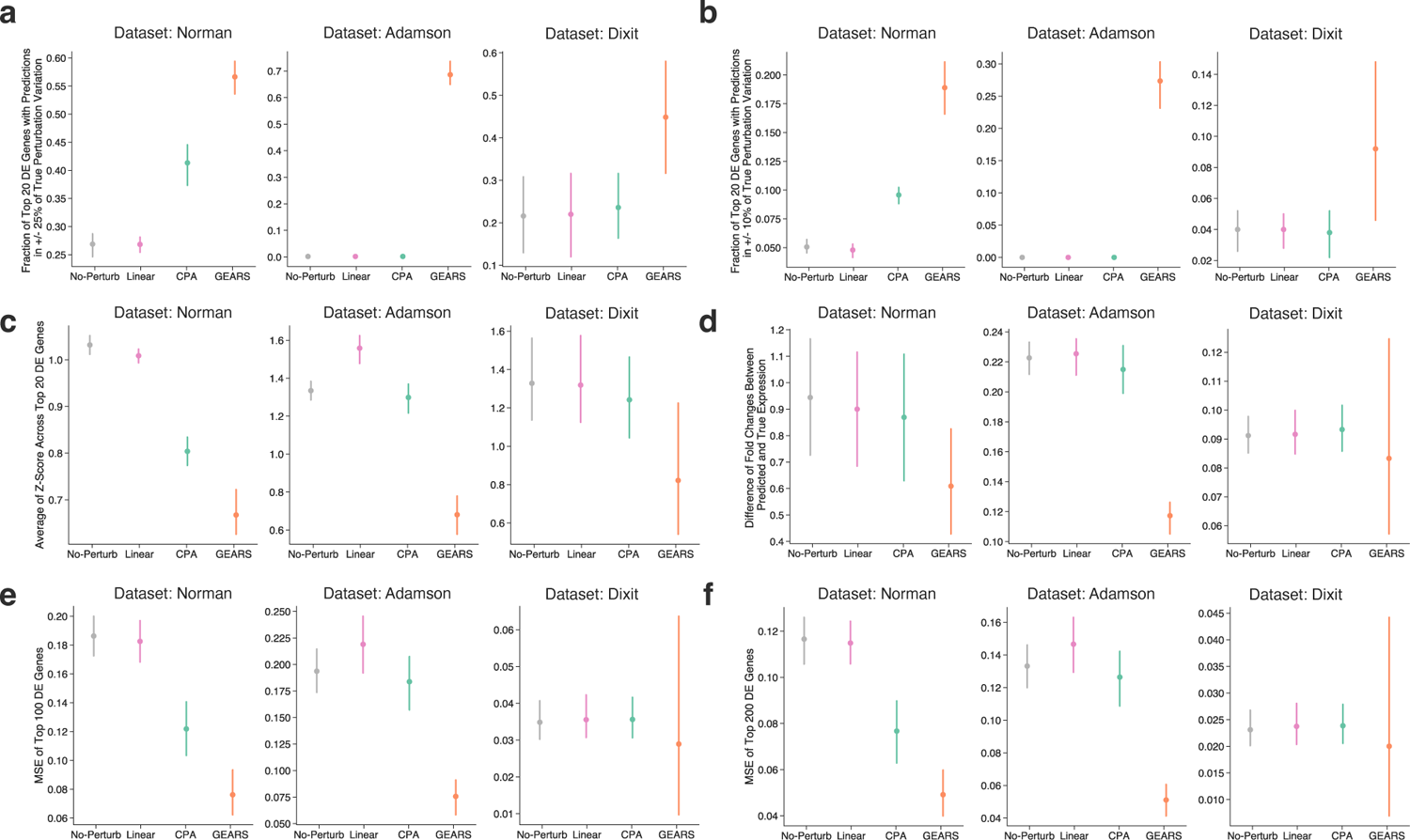
Comprehensive evaluation establishes robustness of GEARS’s prediction of post-perturbation expression. **(a)** Fraction of the top 20 differentially expressed genes for each perturbation that have predicted post-perturbation expression within the 40^*th*^ percentile and the 60^*th*^ percentile of the true post-perturbation expression. **(b)** Fraction of the 20 most differentially expressed genes for each perturbation that have predicted post-perturbation expression within +/- 25% of true post-perturbation expression variation. This corresponds to the interval between the 25^*th*^ percentile and the 75^*th*^ percentile of the true post-perturbation expression. **(c)** Measuring variability in predictions using the average Z-Score across top 20 differentially expressed genes. Z-score was computed using the mean and standard deviation of the true post-perturbation expression distribution for each gene after each peturbation. **(d)** Fold change between predicted post-perturbation expression and true expression. **(e)** MSE in predicted post-perturbation expression for top 100 differentially expressed genes. **(f)** MSE in predicted post-perturbation expression for top 200 differentially expressed genes.

**Extended Data Fig. 2:**
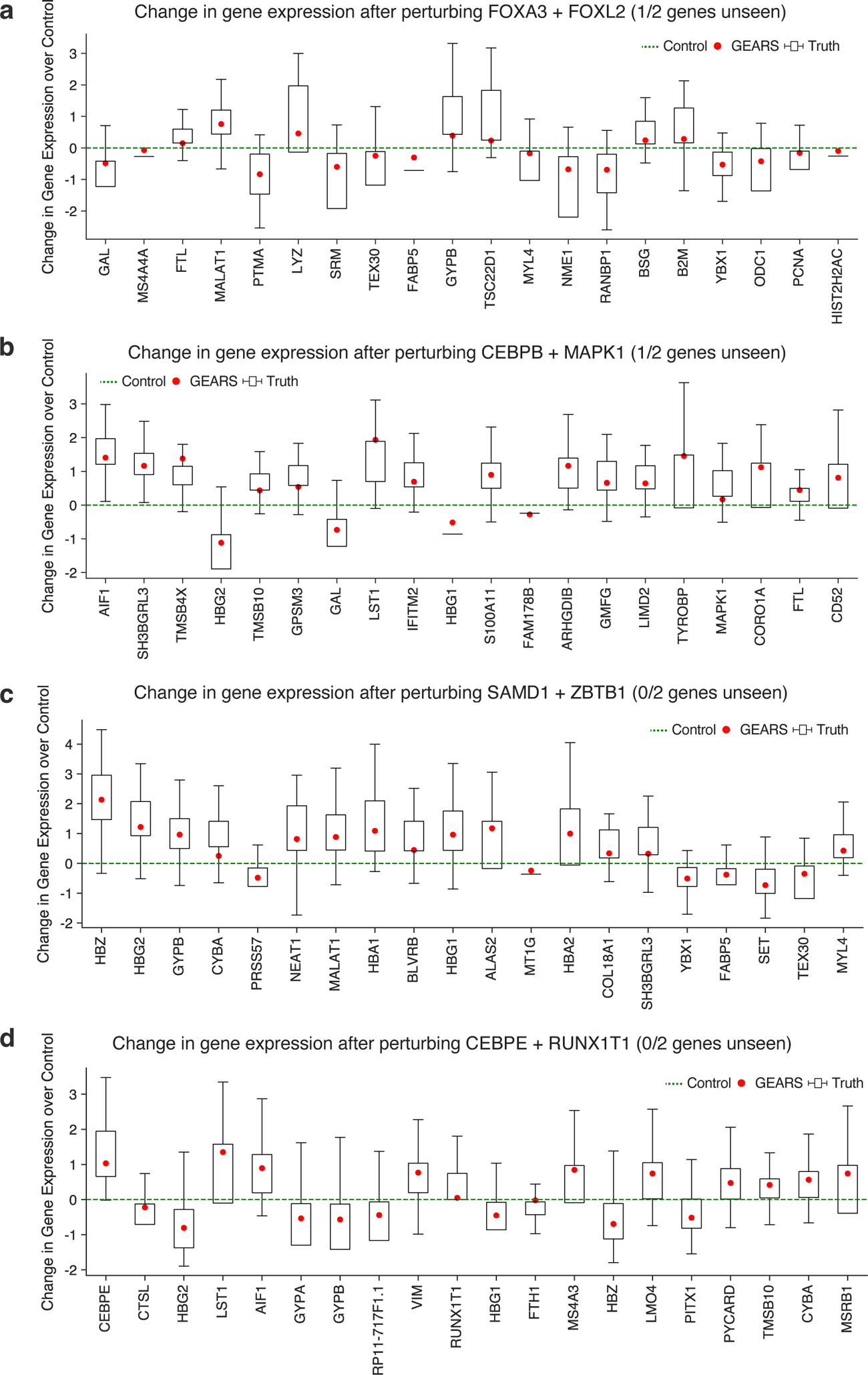
Examples of predicted gene expression across 20 most differentially expressed genes after combinatorial perturbation. **(a)** Change in gene expression after perturbing FOXA3+FOXL2. **(b)** Change in gene expression after perturbing CEBPB+MAPK1. **(c)** Change in gene expression after perturbing FEV+MAP7D1. **(d)** Change in gene expression after perturbing SAMD1+ZBTB1.**(e)** Change in gene expression after perturbing ETS2+IKZF3.**(f)** Change in gene expression after perturbing CEBPE+RUNX1T1.

**Extended Data Fig. 3:**
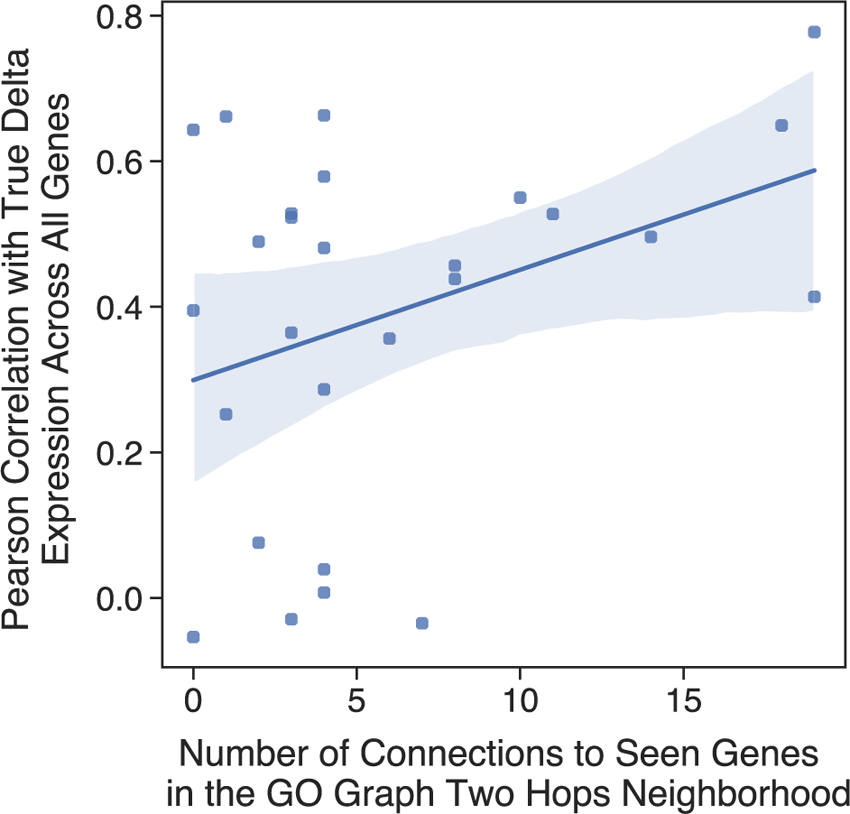
Model performance relationship with network connectivity. Each point in the scatter plot corresponds to a prediction made for a novel single-gene perturbation not seen at the time of training. The y-axis plots the pearson correlation between the true mean post-perturbation differential expression over unperturbed control and the same predicted by GEARS. The x-axis measures the number of connections between the novel perturbed gene and other genes in the network that had been seen at the time of training.

**Extended Data Fig. 4:**
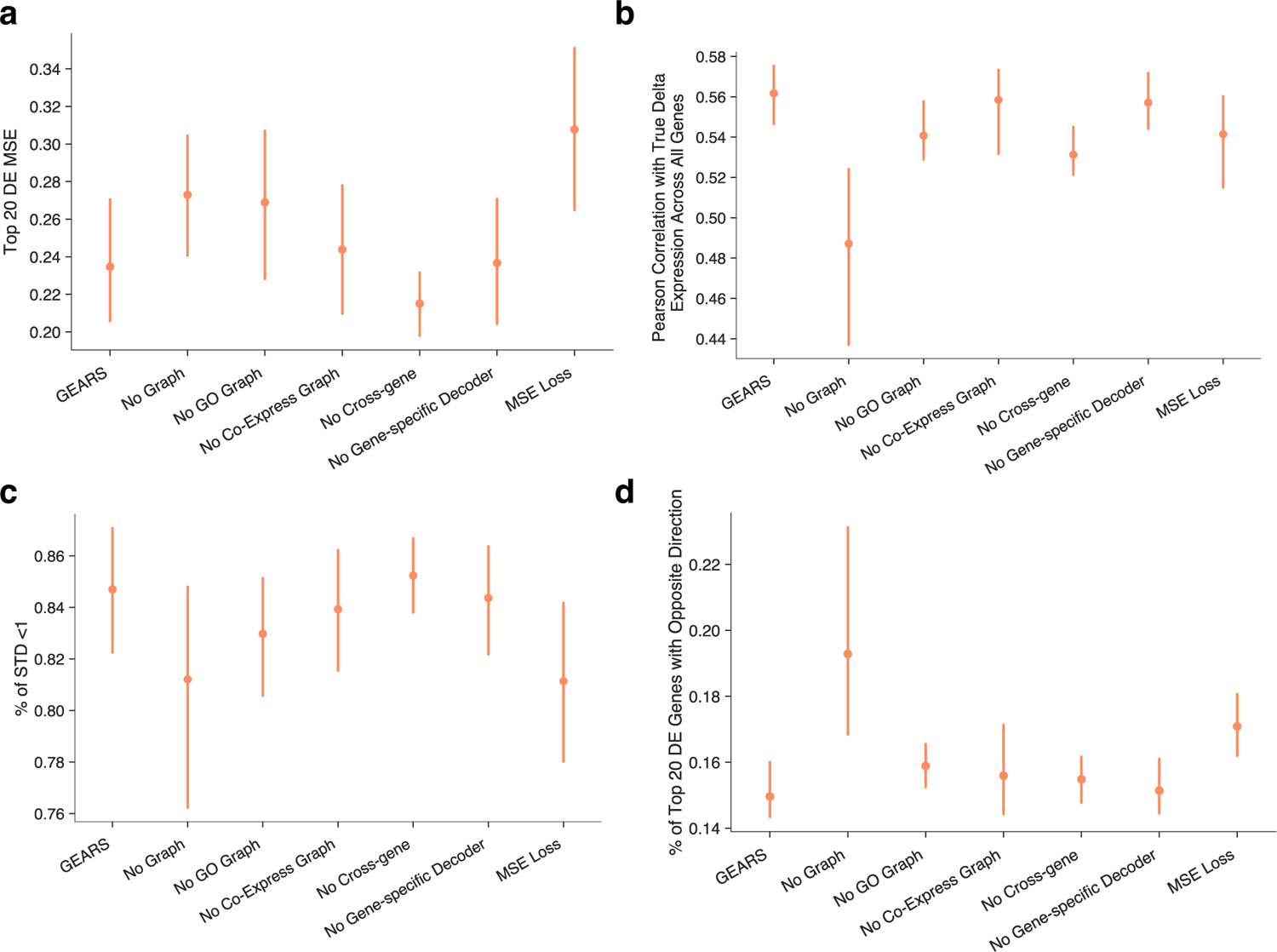
Model Ablation performance. Evaluation of importance of each component of GEARS by testing the performance after removing individual component. “No Graph” removes both the gene ontology graph and co-expression graph; “No GO Graph” removes the gene ontology graph; “No Co-Express Graph” removes the co-expression graph; “No Cross-gene” removes the cross-gene MLP layer; “No Gene-specific Decoder” removes the gene specific decoder MLP and uses a shared MLP instead; “MSE Loss” switches from the auto-focus loss to the regular L2 loss. **(a)** Model ablation in MSE of top 20 most differentially expressed genes. **(b)** Model ablation in pearson correlation between the true mean post-perturbation differential expression over control for across all genes and that which is predicted for the same. **(c)** Percentage of top 20 differentially expressed genes that fall within one standard deviation of the true post-perturbation gene expression distribution. **(d)** Percentage of top 20 differentially expressed genes that have the opposite direction as compared to the true post-perturbation gene expression direction.

**Extended Data Fig. 5:**
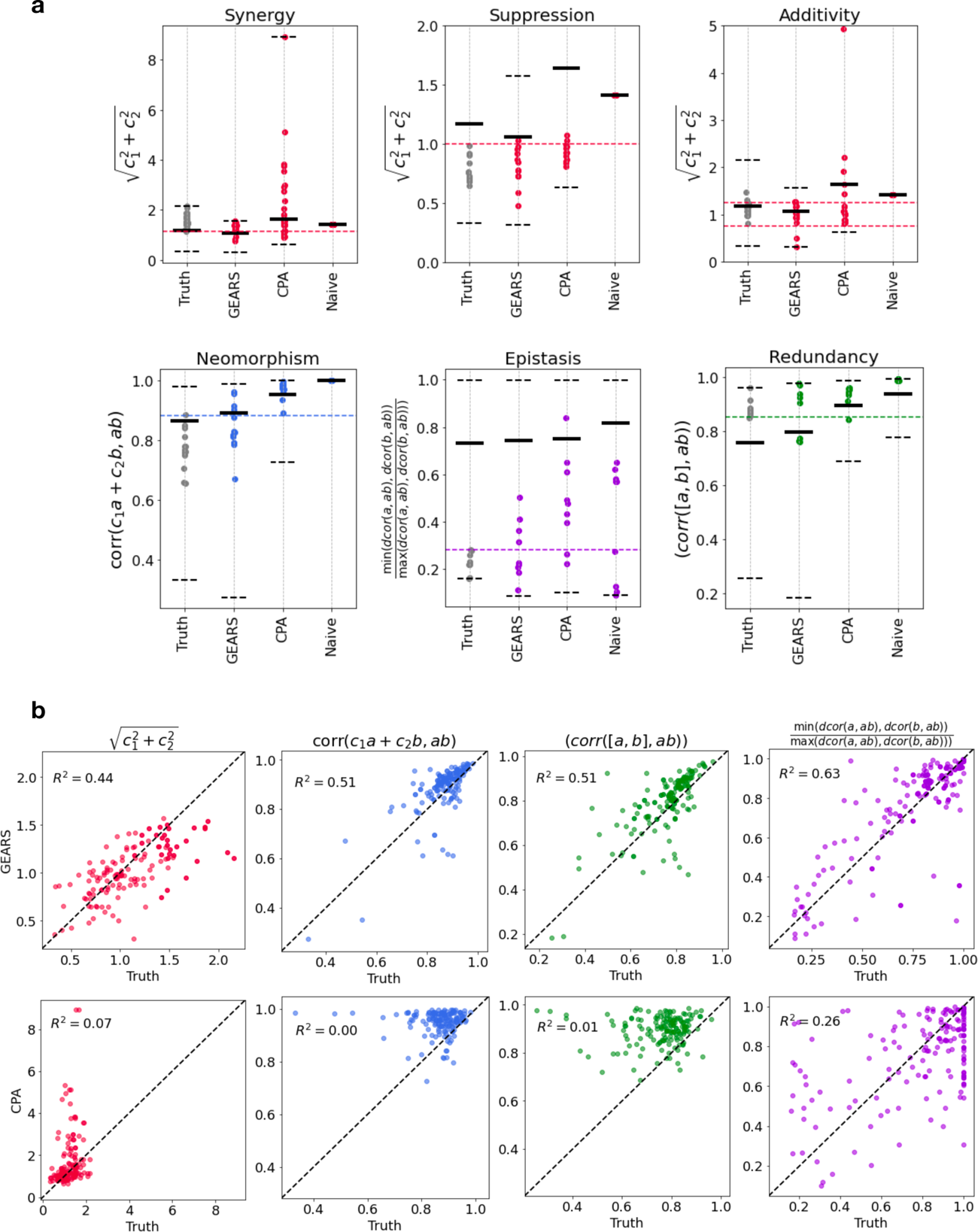
Model performance at predicting GI scores. (a) Each plot in the panel corresponds to predicted or true GI scores for set of combinatorial perturbations that were defined as expressing a specific GI sub-type phenotype in [8]. The gray dots correspond to GI scores computed using true post-perturbation gene expression. The red dots correspond to GI scores computed using predicted post-perturbation gene expression under three different models: GEARS, CPA and Naive models. The naive model here corresponds to a simple additive model where the individual effects of perturbing each of the combining genes are simply added together. The other two models were trained on all the data from [8] while only holding out one specific combinatorial perturbation at a time. Single-gene perturbations for that combination were also seen at the time of training. The metrics on the y-axis correspond to different GI scores and the dotted lines indicate the defined thresholds for determining if a combination is exhibiting a specific GI sub-type phenotype. (b) Scatter plots of GI scores for all 131 2-gene combinatorial perturbations in [8]. The x-axis shows GI scores computed using true post-perturbation gene expression and the y-axis shows scores predicted using predicted post-perturbation gene expression. The top row shows predictions made by GEARS and the bottom row shows predictions made by CPA [26] *R*^2^ refers to the coefficient of determination.

**Extended Data Fig. 6:**
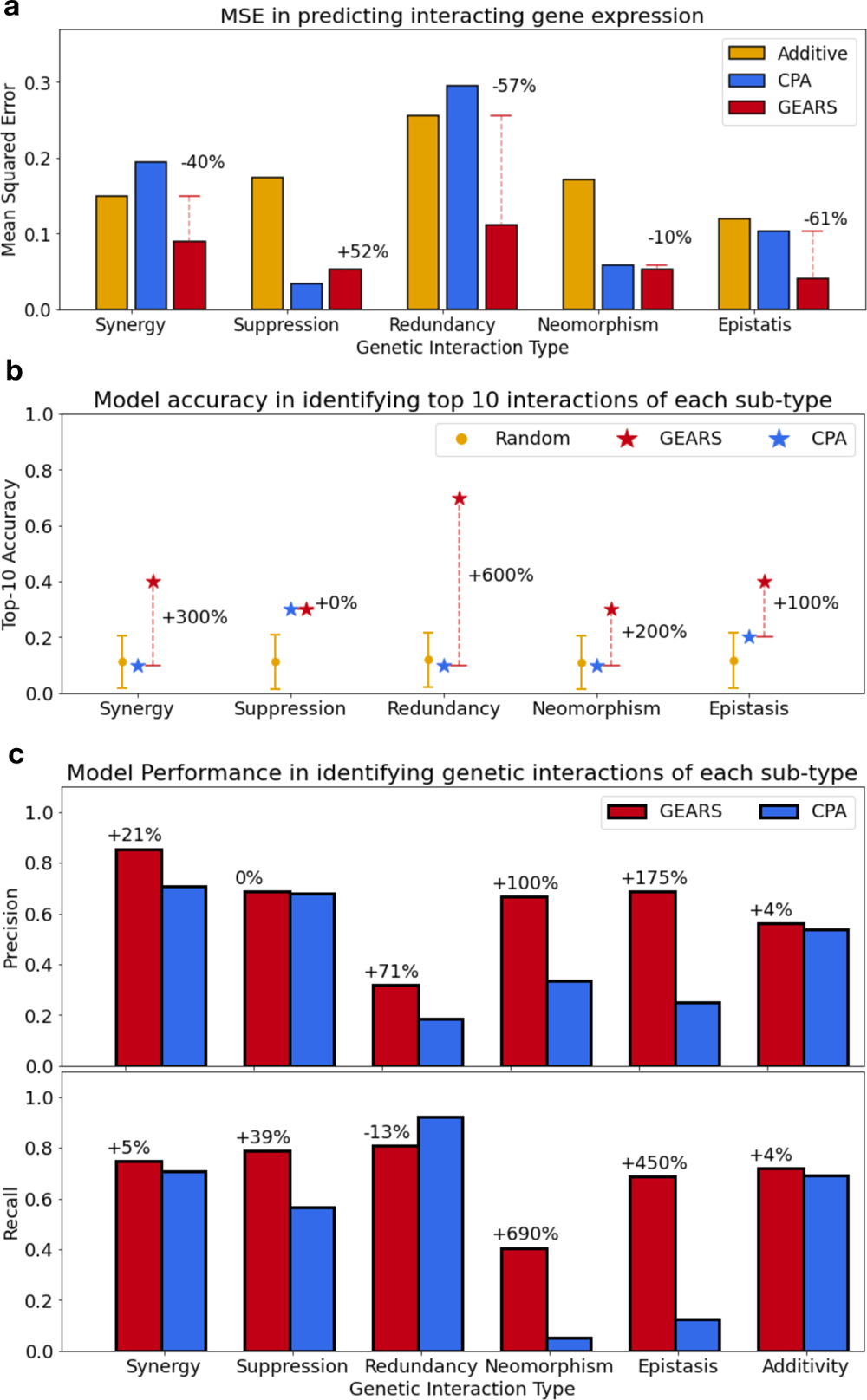
Model performance in predicting genetic interactions. **(a)** Mean Square Error (MSE) in predicting non-additive combinatorial effects between the additive model which assumes that the effect of the combination is just the sum of the two known single-gene perturbation outcomes and GEARS predictions. MSE was measured on the 20 genes with the largest difference between true post-perturbation expression following 2-gene combinatorial perturbation and the additive prediction for that combination. Combinations are categorized on the x-axis by genetic interaction (GI) sub-types defined in [8]. **(b)** Top 10 accuracy in predicting GIs: Model accuracy in predicting the set of 10 strongest interactions for each GI sub-type as determined using true expression. **(c)** Precision and recall in predicting GIs. GIs were identified across all combatorial perturbations using GI sub-type specific thresholds (Methods) applied to model predicted GI scores as well as true GI scores.

**Extended Data Fig. 7:**
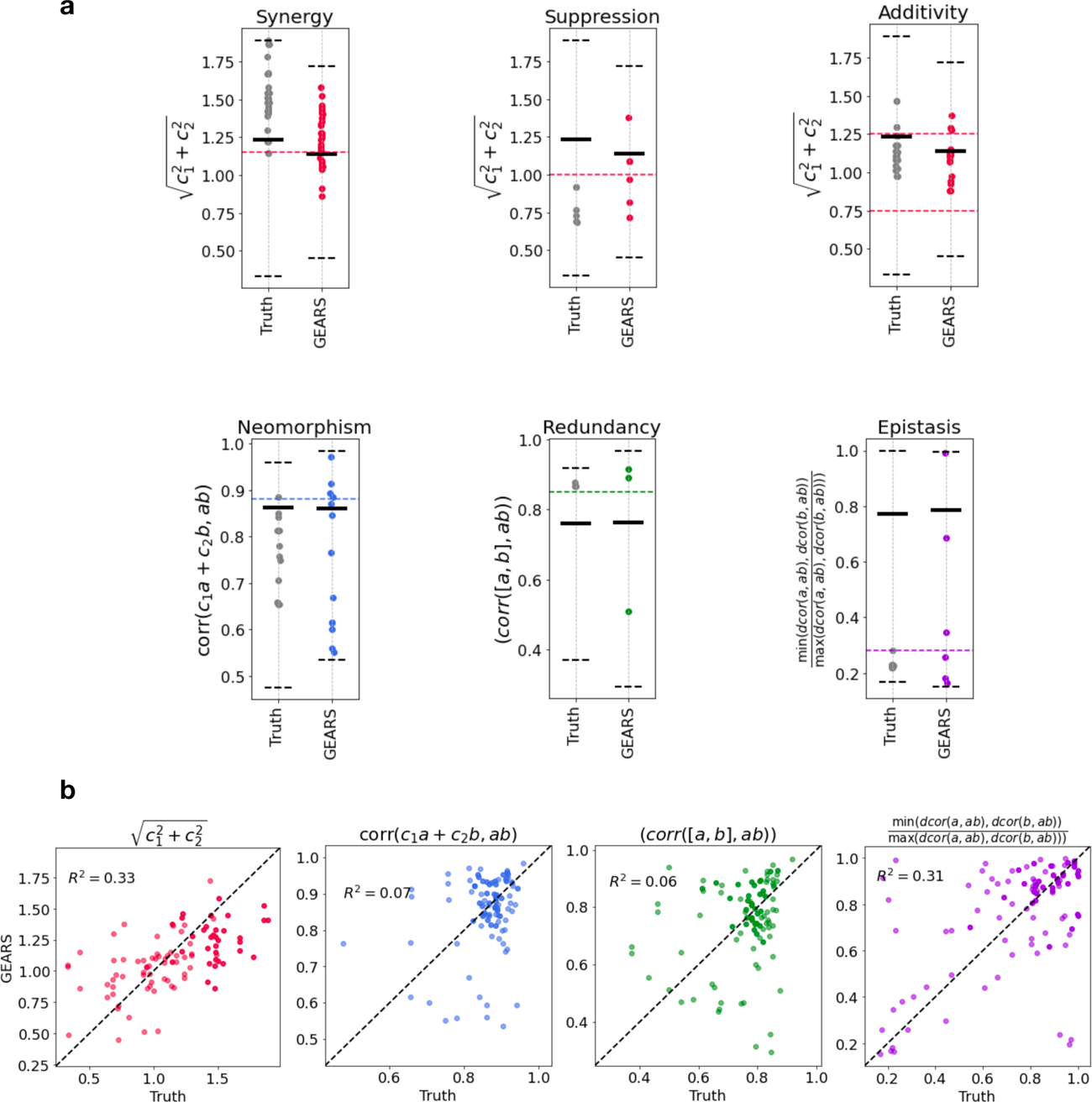
Model performance at predicting GI scores when one of the combining genes is not seen at the time of training. All measurements were performed only on that half of the 262 predicted combinations that had lower uncertainty. Each combination is predicted by the model twice, each time holding out one of the combining genes in the test set. These are treated as distinct predictions. (a) Each plot in the panel corresponds to predicted or true GI scores for set of combinatorial perturbations that were defined as expressing a specific GI sub-type phenotype in [8]. The gray dots correspond to GI scores computed using true post-perturbation gene expression. The red dots correspond to GI scores computed using predicted post-perturbation gene expression from GEARS. GEARS was trained on all the data from [8] while only holding out all combinations that contained one specific gene, making it a novel unseen gene at the time of prediction. The metrics on the y-axis correspond to different GI scores and the dotted lines indicate the defined thresholds for determining if a combination is exhibiting a specific GI sub-type phenotype. (b) Scatter plots of GI scores for all 131 2-gene combinatorial perturbations in [8]. The x-axis shows GI scores computed using true post-perturbation gene expression and the y-axis shows scores predicted using predicted post-perturbation gene expression. The top row shows predictions made by GEARS and the bottom row shows predictions made by CPA [26] *R*^2^ refers to the coefficient of determination.

**Extended Data Fig. 8:**
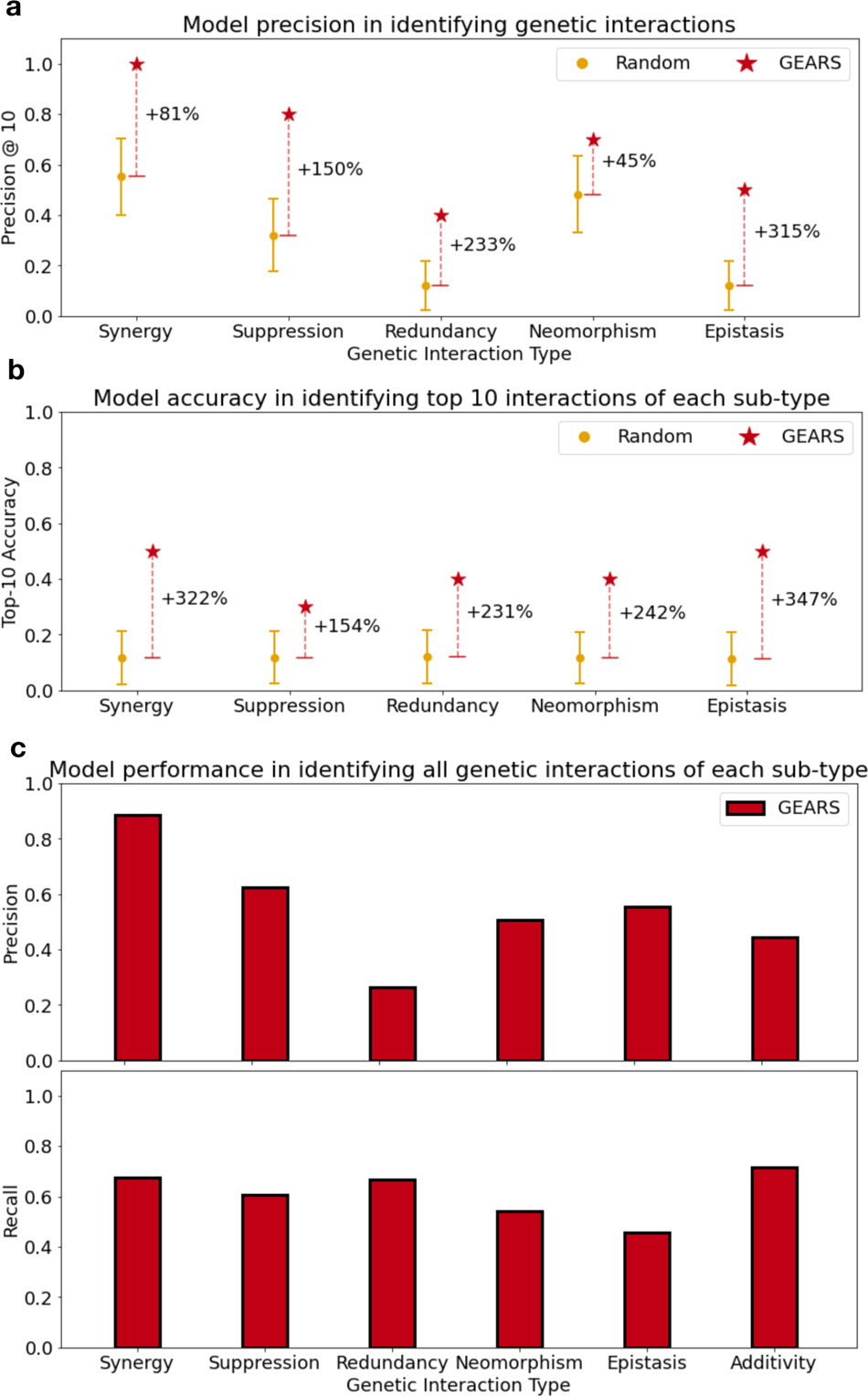
Model performance in predicting genetic interactions when one of the interacting genes is not seen perturbed at the time of training. All measurements were performed only on that half of the 262 predicted combinations that had lower uncertainty. Each combination is predicted by the model twice, each time holding out one of the combining genes in the test set. These are treated as distinct predictions. (a) **Precision@10**: Model precision in predicting genetic interactions from 110 (possibly non-unique) 2-gene combinations. The combinations were ranked using the corresponding genetic interaction (GI) scores for each GI sub-type (Methods). Precision@10 was calculated as the fraction of the top 10 combinations predicted by GEARS for each GI sub-type that also showed that GI phenotype based on true post-perturbation expression. (b) **Top 10 Accuracy**: For each GI sub-type, this metric is the size of the intersection between the set of 10 strongest interactions predicted by the model and the 10 strongest interactions determined using true expression. **Precision and Recall**: GIs were identified across all combatorial perturbations using GI sub-type specific thresholds (Methods) applied to model predicted GI scores as well as true GI scores.

**Extended Data Fig. 9:**
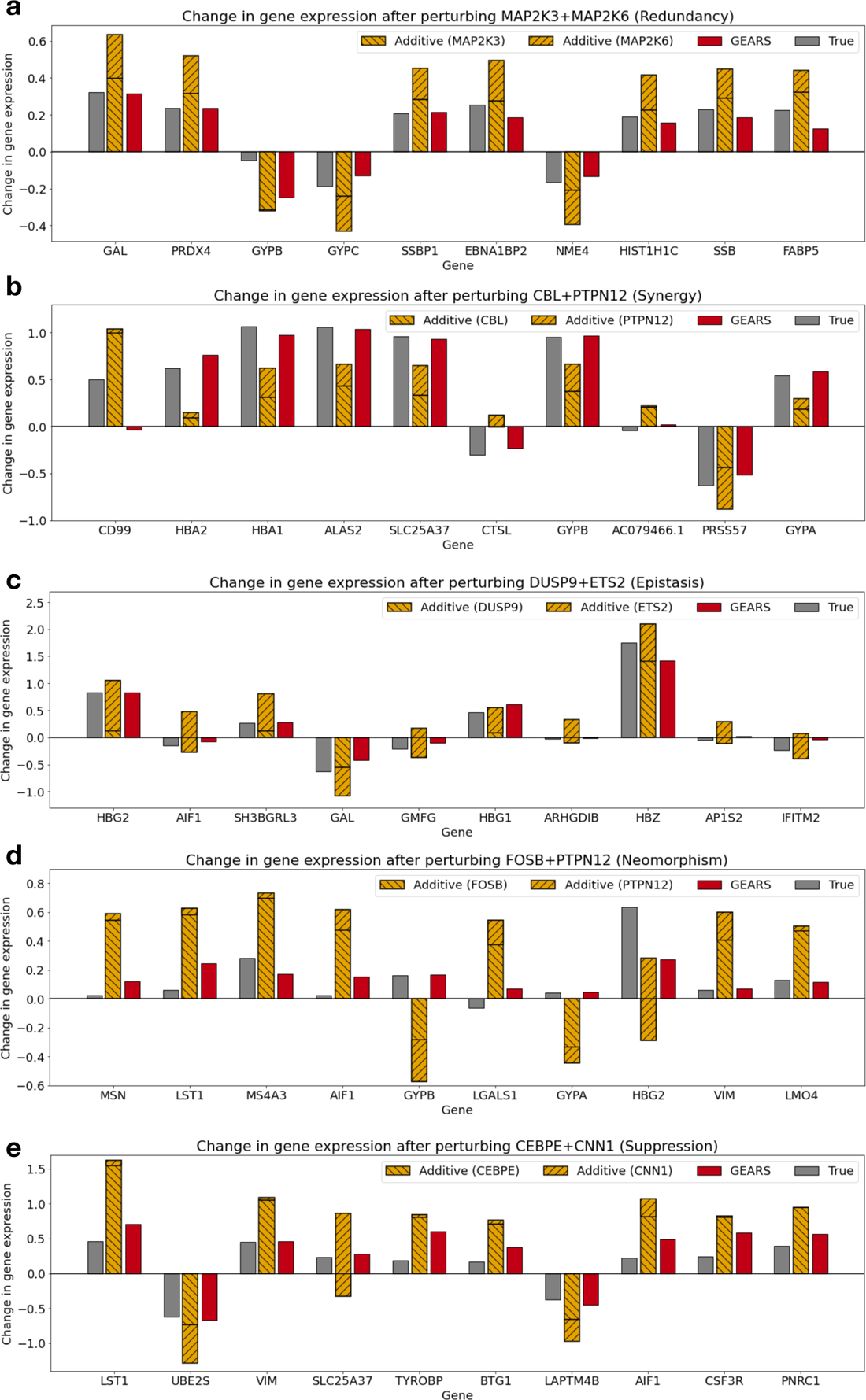
GEARS predicts non-additive combinatorial effects across all GI sub-types. Each panels shows a change in gene expression over unperturbed control after perturbing a combination of genes corresponding to a specific GI sub-type. The gray bars show the true post-perturbation gene expression change over unperturbed control for a particular gene. The hatched yellow bars show the true post-perturbation gene expression for each of the two single-gene perturbations performed individually. The naive additive model assumes that the effect of the combination is just the sum of the two known single-gene perturbation outcomes. The red bar indicates the prediction made by GEARS. The genes on the x-axis are those with the largest difference between true post-perturbation expression following combinatorial perturbation and the additive prediction for that combination. The different GI sub-types considered are: (a) Redundancy (b) Synergy (c) Epistasis (d) Neomorphism (e) Suppression

**Extended Data Fig. 10:**
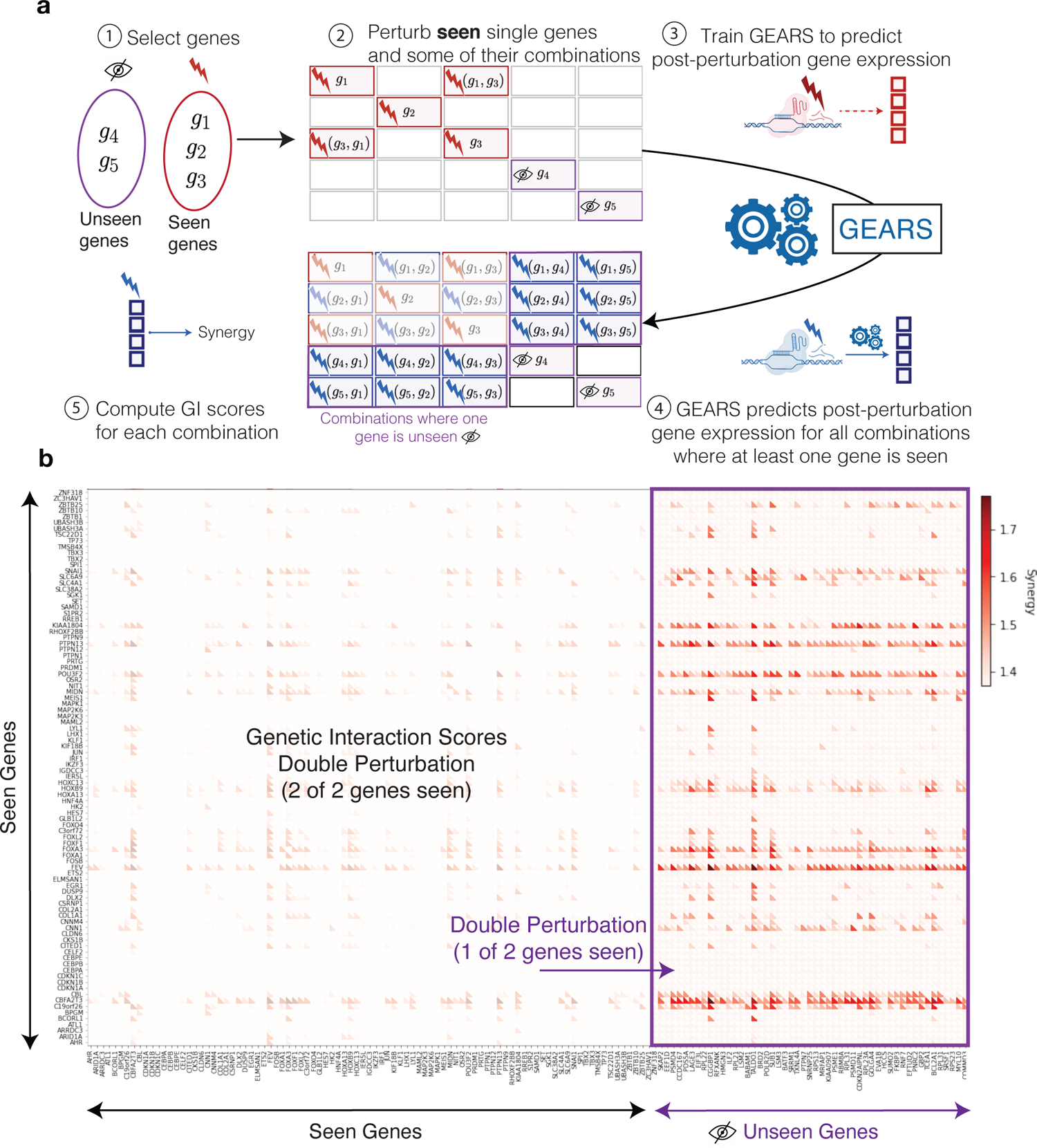
Model predicted genetic interactions for all combinations where at most one of the combining genes is not seen at the time of training. **(a)** Workflow for predicting all pairwise genetic interactions for a set of genes where a subset of those genes are not seen perturbed individually at the time of training. (1) Given a set of seen and unseen genes, (2) the first step is to experimentally perturb all single seen genes and measure the post-perturbation gene expression. The same experiment can also be performed on a selection of combinations depending upon time and cost. (3) GEARS is then trained using this data to predict post-perturbation gene expression. (4) After training, GEARS predicts post-perturbation gene expression for all pairwise combinations of seen and unseen genes where at least one gene has been seen perturbed individually at the time of training. (5) Synergy GI score for each combination can then be calculated using post-perturbation gene expression. **(b)** GEARS predicted genetic interaction scores for synergy for all combinations of genes in the seen and unseen gene sets where at least one gene has been seen perturbed individually at the time of training.

