## Supplementary Information for "GEARS: Predicting transcriptional outcomes of novel multi-gene perturbations"

#### Generating UMAPs and clustering of post-perturbation gene expression data.

The UMAPs in Figure 3(b) and figure 3(c) were generated using GEARS-predicted post-perturbation gene expression profiles of all pairwise combinations of the 105 1-gene perturbations from the the Norman et al. [8] dataset. Figure 3(c) shows this complete UMAP with all possible 1-gene and 2-gene combinations for this dataset. Figure 3(b) was plotted using GEARS-predicted post-perturbation gene expression profiles of only those perturbations from [8] (105 1-gene perturbations and 131 2-gene perturbations) used to train GEARS. Thus, Figure 3(b) is a subset of the data in Figure 3(c). The UMAP manifold was computed only once for both figures to enable a direct comparison.

Clustering was performed using Leiden clustering with default parameters set in scanpy (resolution = 1) (Supplementary Figure 4). Clusters shown in Figure 3(b) and 3(c) were labelled using phenotypic labels from Norman et al. [8]. If any cluster or set of clusters contained perturbations that were labelled as exhibiting a specific phenotype in Norman et al., then that whole cluster or set of clusters was labelled as showing that specific phenotype in Figure 3(b) or 3(c). All other clusters were not given phenotypic labels in these figures.

**Differential expression analysis for novel phenotypic cluster identified by GEARS.**

The novel phenotypic cluster identified by GEARS showed high expression of known erythroid lineage-specific marker genes. *HBG1, HGB2, HBZ, HGA1, HBA2, GYPA, ERMAP* were identified as erythroid lineage-specific marker genes in [8]. Out of these, *HBG1, HBZ, HBA2, GYPA* were present among the genes whose expression GEARS was trained to predict (Methods). We see a higher expression of all these four marker genes in the new phenotypic cluster than in any experimentally tested perturbation (Supplementary Figure 3)

For further validation of the novel phenotypic cluster discovered by GEARS we used single-cell gene expression data from Tabula Sapiens. We chose to use the data from the bone marrow organ as it contains hematopoetic progenitor cells, which is the cell type most similar to the K562 lymphoblasts that were used for perturbational experiments in Norman et al. [8]. The bone marrow dataset also contains all the cell types that hematopoetic progenitor cells can differentiate into. We identified proerythroblasts as a cell type that represents an early stage in the erythroid lineage. The differential expression (DE) between hematopoetic progenitor cells and proerythroblasts was use to represent the transition to an early stage in the erythroid lineage in Tabula Sapiens. Differentially expressed genes were identified using scanpy and the log fold change in expression was computed using diffxpy. Any gene with absolute value of log fold change in expression greater than 10 or less than 0.01 was not considered. A vector $v_e = [\Delta g_1, \Delta g_2 \ldots, \Delta g_k]^T$ consisting of the log fold change in expression for all $k$ differentially expressed genes was constructed for the transition from hematopoetic progenitor cells to proerythroblasts. A similar DE vector $v_p$ for the same $k$ genes was constructed for each of the perturbations that GEARS predicted the outcome for. A dot product $v_e \cdot v_p$ was computed between the DE vector for each perturbation and the DE vector for the transition to proerythroblasts (Figure 3). We see that the new phenotypic cluster exhibits a novel phenotype that is most similar to the transition to proerythroblasts in Tabula Sapiens. Hence, is it both novel and biologically meaningful.

### Supplementary Figures

- **Supplementary Information Fig. 1**: Data split matrix
- **Supplementary Information Fig. 2**: Subgroup analysis of additional evaluation metrics for predicting post-perturbation expression
- **Supplementary Information Fig. 3**: Erythroid-lineage marker expression for GEARS predicted perturbation outcomes
- **Supplementary Information Fig. 4**: Leiden clustering of post-perturbation gene expression profiles for each perturbation predicted by GEARS
- **Supplementary Information Fig. 5**: Variation in model performance between predictions with low uncertainty and others.

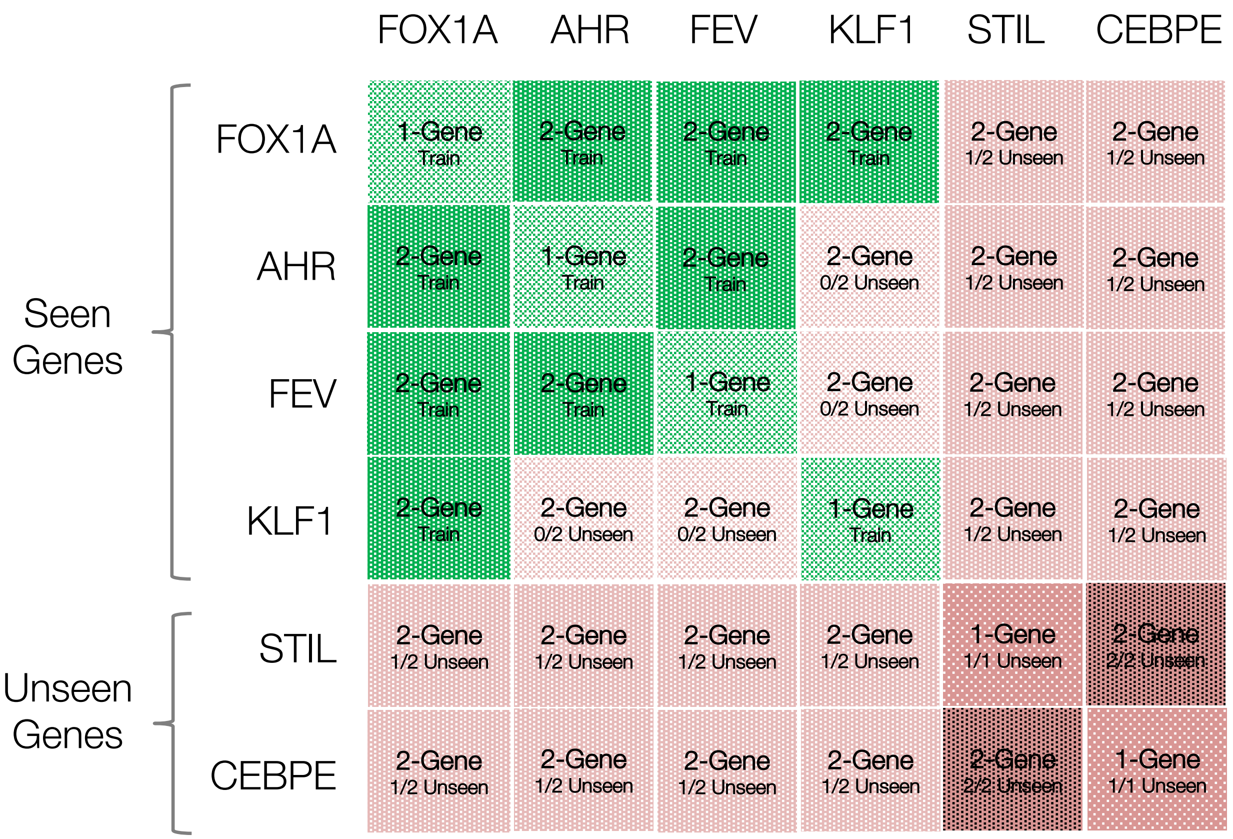


**Supplementary Information Fig. 1: Data split matrix** A sample data split illustration to describe how the different perturbation categories were defined on the basis of training set composition. Genes that were *seen* experimentally perturbed in the training data and *unseen* genes are marked on the vertical axis. 1-gene perturbations of seen genes were included in the training set (**1-Gene, Train**). 1-gene perturbations of unseen genes were included in the test set (**1-Gene, 1/1 Unseen**). 2-gene combinatorial perturbations with one gene unseen were included in the test set (**2-Gene, 1/2 Unseen**) as were those with two genes unseen (**2-Gene, 0/2 Unseen**). 2-gene combinatorial perturbations with both genes seen were randomly split between train (**2-Gene, Train**) and test (**2-Gene, 2/2 Unseen**).

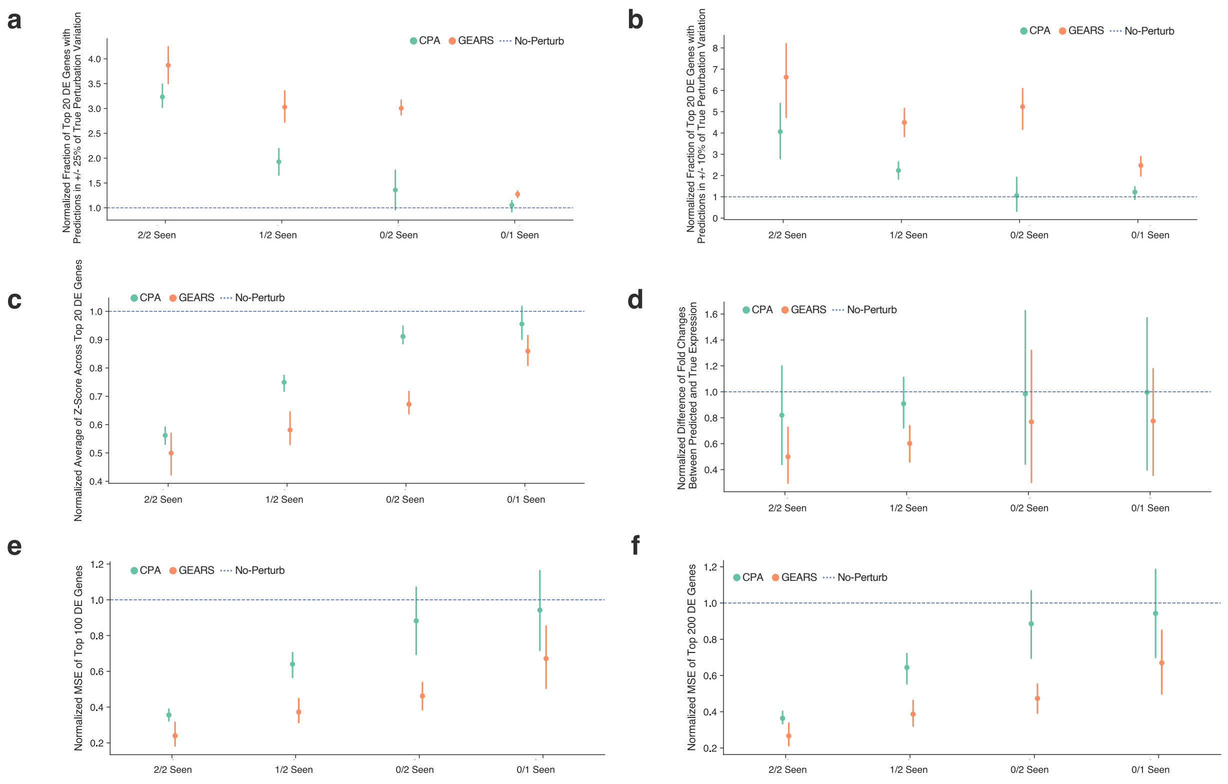


**Supplementary Information Fig. 2: Subgroup analysis of additional evaluation metrics for predicting post-perturbation expression.** **(a)** Fraction of top 20 differentially expressed genes with predictions fall in +/- 10% of true perturbation variation. **(b)** Fraction of top 20 differentially expressed genes with predictions fall in +/- 25% of true perturbation variation. **(c)** Average of Z-Score across top 20 differentially expressed genes. **(d)** Difference of fold changes between predicted and true expression. **(e)** MSE of Top 100 differentially expressed genes. **(f)** MSE of Top 200 differentially expressed genes.

a

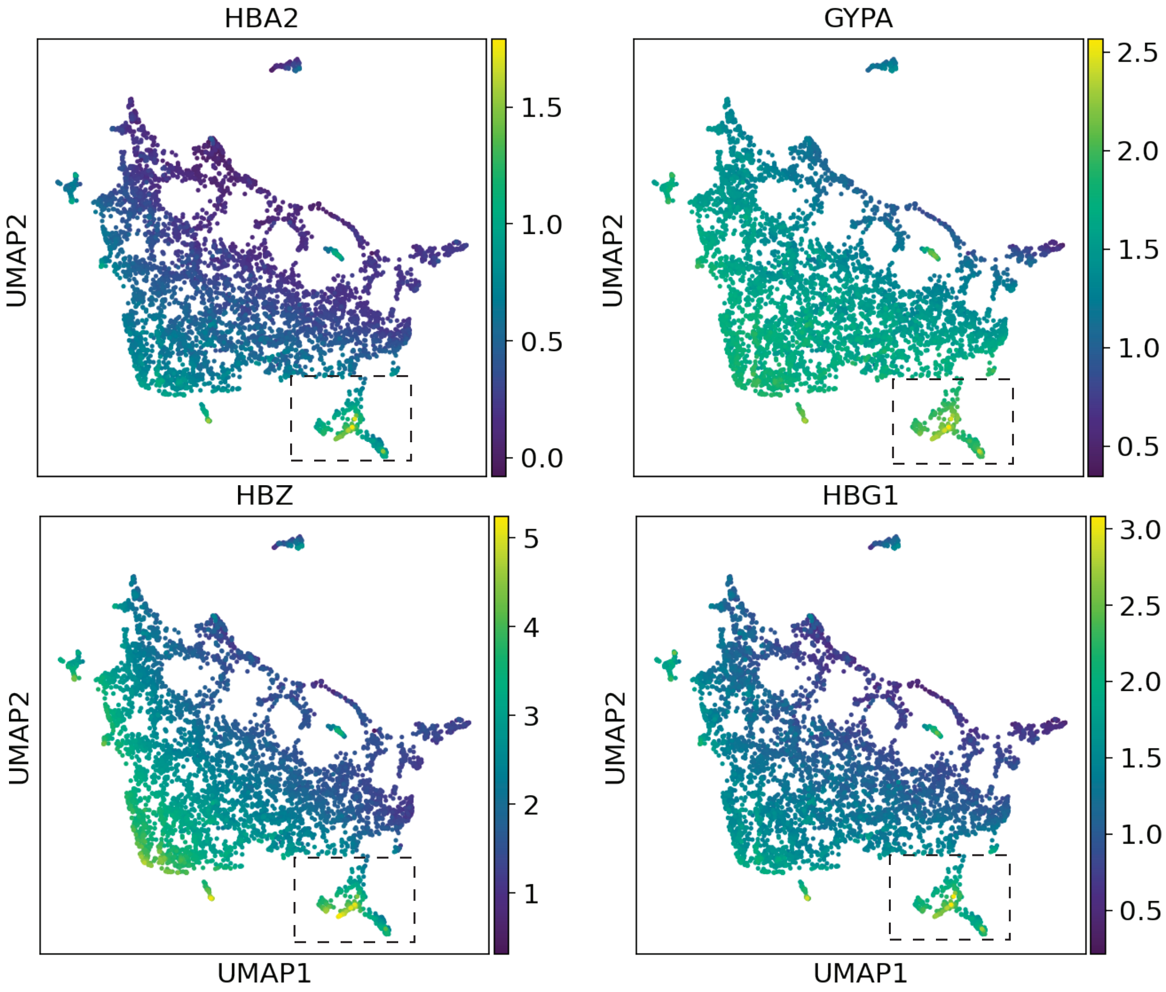


**Supplementary Information Fig. 3: Erythroid-lineage marker expression for GEARS predicted perturbation outcomes**

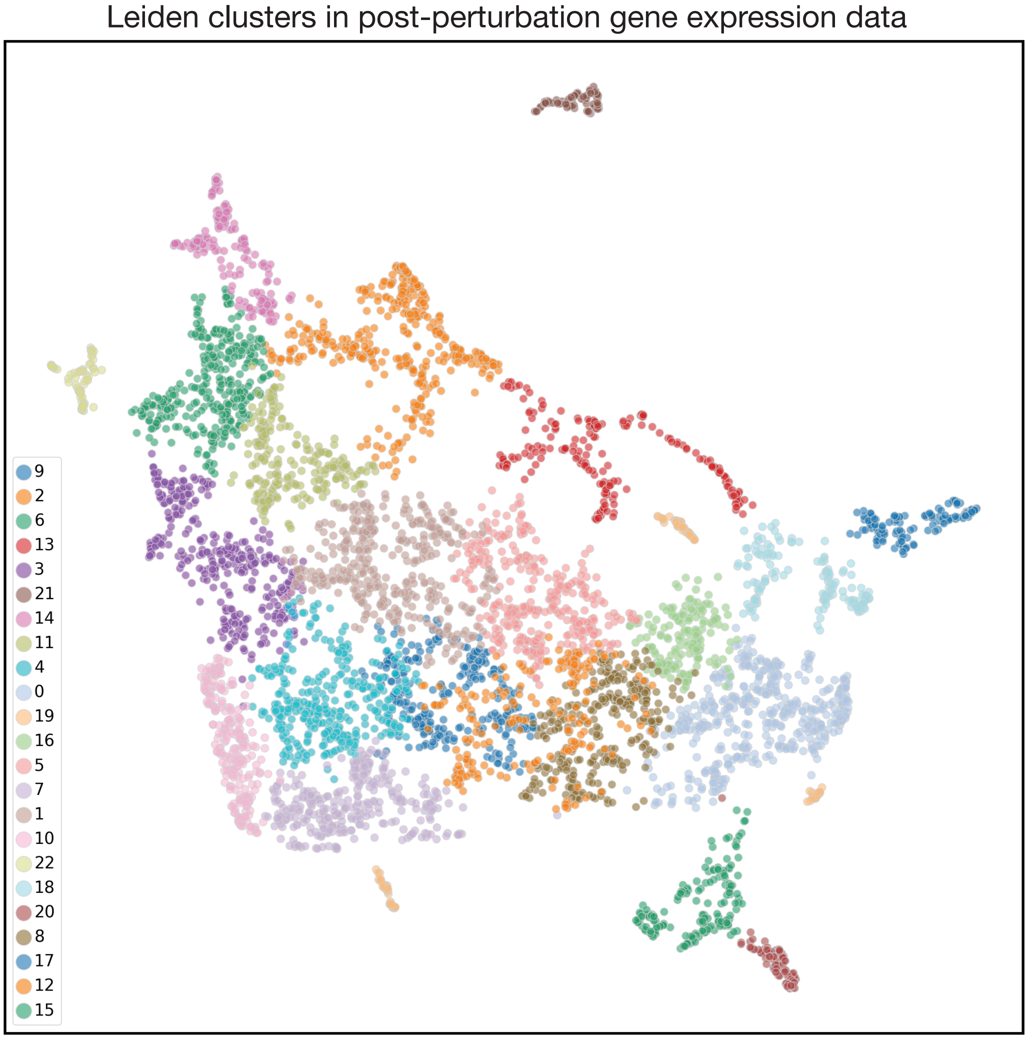


| Phenotype | Clusters |
|---|---|
| Pro Growth | 2, 4, 16, 22 |
| Pioneer Factors | 0 |
| Granulocyte Marker | 17 |
| Megakaryocyte Marker | 13 |
| Erythroid Marker | 10, 7 |
| High Erythroid Marker Expression (New) | 15 |

**Supplementary Information Fig. 4: Leiden clustering of post-perturbation gene expression profiles for each perturbation predicted by GEARS**. Includes all 105 1-gene perturbations and 131 2-gene perturbations that GEARS was trained on. Table shows the assignment of clusters to phenotypic labels based on how perturbations were labelled in [8]. Any cluster containing a perturbations assigned a phenotypic label in [8] was labelled as exhibiting that phenotype in Figure 3(b) and 3(c)

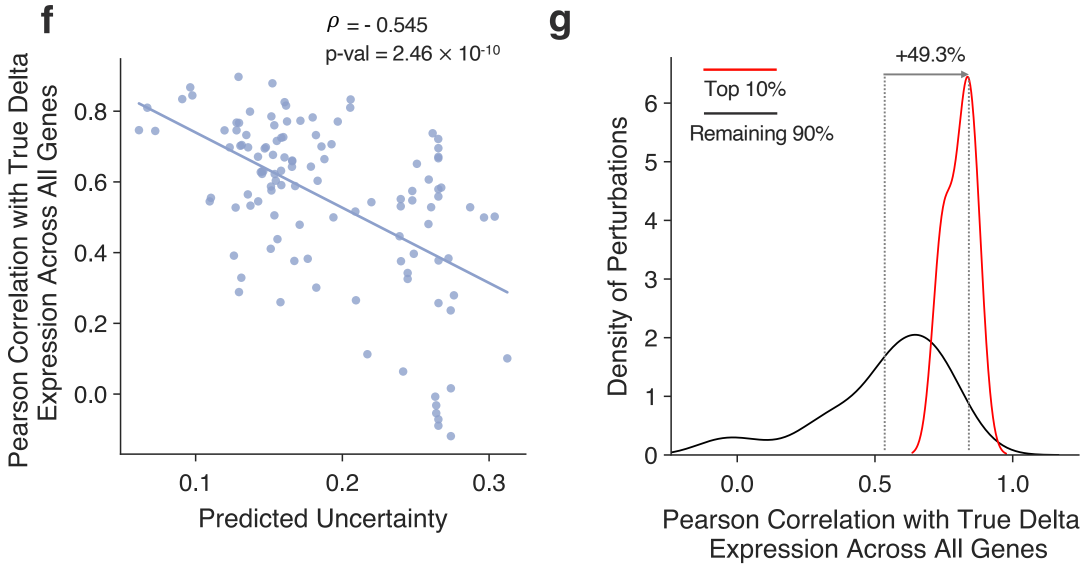


**Supplementary Information Fig. 5: Variation in model performance between predictions with low uncertainty and others**. The x-axis measures the Pearson correlation of predicted post-perturbation differential expression values over control and true post-perturbation differential expression over control for all genes. The red distribution corresponds to perturbations with the lowest 10% predicted uncertainty while the black curve corresponds to all other perturbations
